# Mutations in severe human H5N1 cases facilitate evasion from human mucus and antivirals

**DOI:** 10.64898/2026.07.28.740943

**Authors:** Ksenia Sukhova, Jiayun Yang, Antonio Di Maio, Jin Yu, Maria R. Oliveira, Hannah Hens, Jean-Remy Sadeyen, Sareeta Bagri, Sam Jones, Jack A. Hassard, Joelle Mettier, Jie Zhou, Apoorva D. Srivastava, Naina Mathur, Benjamin Schumann, Stuart M. Haslam, Munir Iqbal, Yan Liu, Thomas P. Peacock, Wendy S. Barclay, Hannah Klim

**Affiliations:** Department of Infectious Disease, Imperial College London, Exhibition Rd, London, United Kingdom, SW7 2AZ; The Pirbright Institute, Ash Rd, Woking, United Kingdom GU24 0NF; Department of Metabolism, Digestion and Reproduction, Imperial College London, Hammersmith Hospital, Du Cane Road, London, United Kingdom, W12 0NN; Department of Life Sciences, Imperial College London, Exhibition Rd, London, United Kingdom, SW7 2AZ; Department of Chemistry, Imperial College London, London, United Kingdom, W12 0BZ; Chemical Glycobiology Laboratory, The Francis Crick Institute, London, United Kingdom, NW1 1AT; Faculty of Chemistry and Food Chemistry, TUD Dresden University of Technology, 01062 Dresden, Germany

## Abstract

In late 2024, two individuals in Canada and the United States were treated in intensive care for acute respiratory distress caused by infection with the avian Influenza A Virus H5N1 2.3.4.4b genotype D1.1. Viral sequence data obtained from sampling these patients indicated mixed alleles at haemagglutinin (HA) positions 190 and 226. Mutations at these positions are key determinants of HA usage of α2,6-linked sialic acids (SA), the most abundant influenza receptors in human upper respiratory tracts. Thus, these mutations raised concerns about human adaptation and pandemic potential of the H5N1 virus. In this study, we investigated the impact of the mutations at residues 190 and 226 in H5 HA. We studied the receptor binding properties, cell entry phenotypes and fitness impacts of the mutations using recombinant proteins, pseudotyped lentiviruses, and in the context of influenza viruses using reverse genetics. The mutations did not confer any detectable α2,6-linked sialic acid receptor usage either alone or in combination. Rather, viruses carrying these mutations exhibit weakened binding towards α2,3-linked sialic acid receptors. This correlated with an enhanced capacity to evade human airway mucus, and a reduced susceptibility to oseltamivir and zanamivir. This research underscores that in addition to the way HA interacts with SA as entry receptors, other factors that impact the HA/NA balance might influence the evolutionary trajectory of a zoonotic virus in the human respiratory tract. This study presents a new paradigm for the evolutionary drivers of HA, where reduced sialic acid binding can serve as an advantage for escape from host barriers and antivirals.

## Introduction

Since 2020, highly pathogenic avian influenza A viruses (IAV) from the H5N1 clade 2.3.4.4b have spread rapidly among avian species across six continents^1^ and infected at least 13 mammalian host species^2–13^. During this outbreak, there have been alarming numbers of cases in humans. As part of the ongoing H5N1 2.3.4.4b outbreak in the United States since 2024, there have been 70 confirmed human cases to date linked primarily to infected animals in dairy herds (41) and poultry farms (24), with 5 cases linked to other animals or unknown sources^14^. These cases have mostly been mild, with common symptoms including conjunctivitis and fever and no evidence of human-to-human transmission^15,16^. However, two severe human cases of H5N1 2.3.4.4b identified in North America in late 2024 are of particular concern because of genetic mutations detected in the most abundant viral surface protein and immunogen: haemagglutinin (HA).

In November 2024, a 13-year-old in British Columbia, Canada was treated in intensive care for acute respiratory distress caused by infection with the H5 2.3.4.4b genotype D1.1^17^. Viral sequence data obtained from samples from this patient indicated mixed alleles at residues 190 and 226 in mature H3 numbering (186 and 222 in mature H5 numbering, respectively). At residue 190 there was a mixture of 72% glutamic acid (E) and 28% aspartic acid (D), while at residue 226, 65% glutamine (Q) and 35% histidine (H)^18^. Due to short sequence reads it was not clear whether the two mutations had linkage or not (e.g. occurring in the same HA protein copy). Similarly, in December 2024, a patient over age 65 was hospitalised and later succumbed to infection with the H5N1 2.3.4.4b genotype D1.1 in Louisiana, USA. This case marked the first detected human death caused by H5N1 in the United States. Sequence analysis of the HA gene in samples isolated from this patient reported mixed alleles at position 190 (92% E, 8% D), along with mixed nucleotides at 138 (88% alanine, 12% valine; residue 134 in H5 numbering) and 186 (65% asparagine, 35% lysine; 182 in H5 numbering)^19^.

Avian influenza viruses HAs, including H5N1 viruses of the 2.3.4.4b lineage, generally show a strong binding preference to glycans containing terminal α2,3-linked sialic acids (SA), which is the predominant receptor type in gastrointestinal tracts of the natural waterfowl reservoir. Conversely human seasonal and pandemic influenza viruses show a strong preference for binding to α2,6-linked SAs, which are abundant in human upper respiratory tracts. For a number of different influenza subtypes, including the H1, H2 and H3 subtypes that caused previous human pandemics, mutations at positions 190 (H1 – E190D) or 226 (H2 or H3 – Q226L) have previously been shown to contribute to the switch in HA preference from α2,3-linked SA to α2,6-linked SA, enhancing replication in human cells and facilitating airborne transmission in the ferret model^20–22^. Therefore mutations at these sites in the H5 HA are a concerning sign of potential human adaptation of the virus^23^.

In this study, we investigated the phenotype of the mutations at residues 190 and 226 in the H5 HA gene of the clade 2.3.4.4b genotype D1.1 British Columbia virus. We found no evidence that the mutations conferred α2,6-linked sialic acid receptor usage. Instead, we demonstrate that, while the mutations at 190 and 226 result in reduced binding of α2,3-linked sialic acids, they confer fitness advantages under certain conditions by enhancing escape from human mucus and improving viral replication in the presence of neuraminidase inhibiting antivirals including in primary cultures of human airway epithelial cells. This research highlights that IAVs with low sialic acid affinity are still viable in human airways, and that this low affinity might be a previously overlooked mechanism for overcoming host defences and evading neuraminidase inhibitor antivirals.

## Results

### Mutations E190D and Q226H do not confer a fitness advantage under native conditions

Two minor frequency mutations E190D and Q226H were reported in the H5 HA of an H5N1 virus isolated from a human in Nov. 2024 in British Columbia, Canada^18^ followed by the recurrence of E190D in a human case in Dec. 2024 in Louisiana, USA^19^. We engineered these mutations either together or singly into the HA from A/BC/PHL-2032/2024(H5N1), combined with mutations to replace the polybasic cleavage site with a monobasic cleavage site as previously described^24^. We rescued four recombinant viruses with these HA gene segments in combination with the homologous N1 NA gene segment and the remaining gene segments from the attenuated laboratory strain A/Puerto Rico/8/1934(H1N1)^25^. Upon rescue, HA and NA genes were sequenced to confirm presence of the expected HA mutations (**Figure 1A**). Viruses were titred by plaque assay in Madin-Darby Canine Kidney (MDCK) cells for use in viral replication kinetics studies (**Figure 1A**), and we noted that all mutants displayed a smaller plaque phenotype in MDCK cells than the wildtype virus.

**Figure 1.**
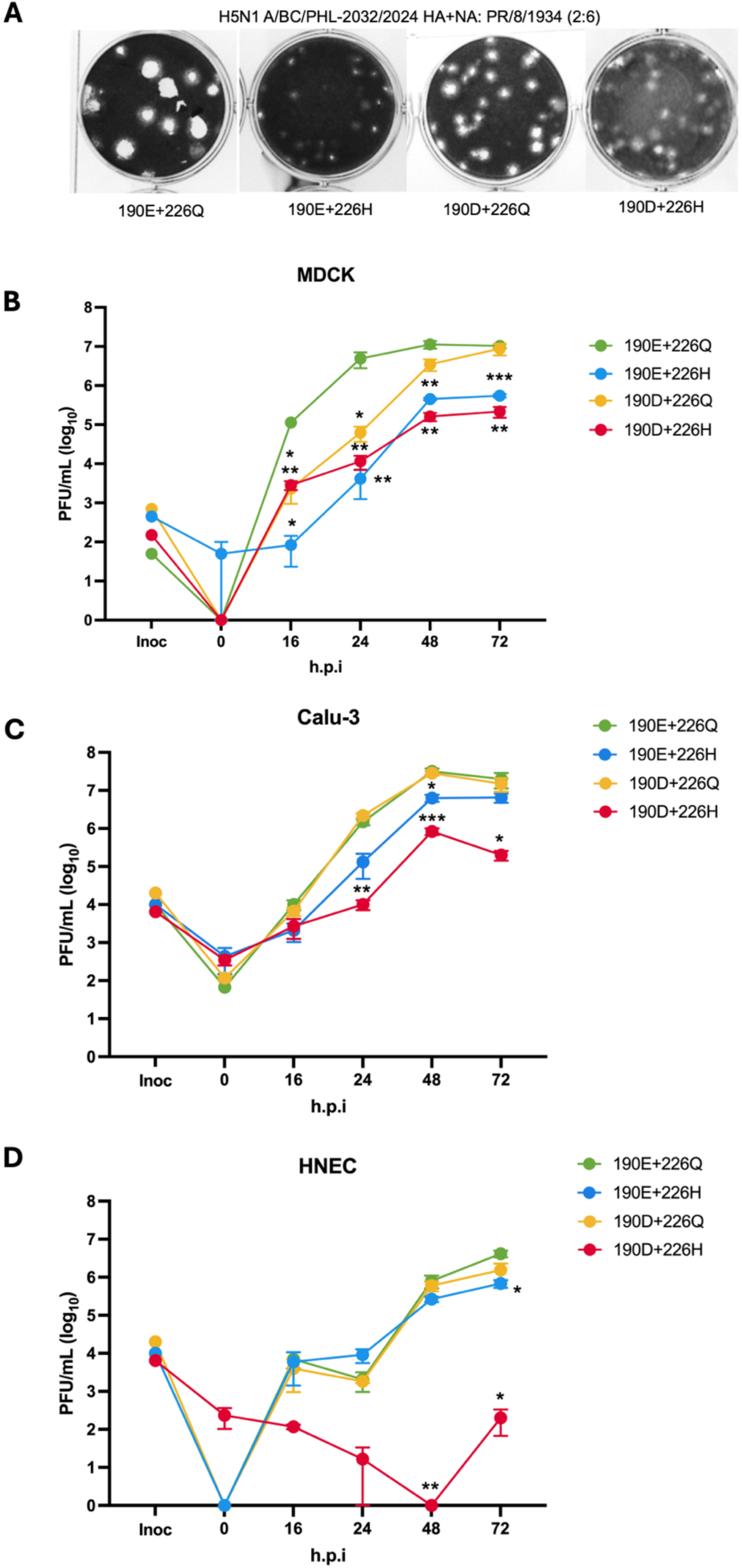
HA mutations E190D and Q226H do not confer an advantage for viral growth kinetics. **A.** Plaque assays of four H5N1 A/BC/PHL-2032/2024 HA+NA: H1N1 A/Puerto Rico/8/1934(2:6) viruses with HA mutations 190E 226Q (avian wildtype), 190E 226H, 190D 226Q, 190D 226H. Viruses were sequenced to confirm correct mutations. **B-D.** Viral growth kinetics of the four H5N1 A/BC/PHL-2032/2024 HA+NA: H1N1 A/Puerto Rico/8/1934(2:6) viruses at a multiplicity of infection (MOI) of 0.01. Supernatant was collected from cells at 0, 16, 24, 48, and 72 hours post infection (h.p.i.). Viral titres were quantified by plaque assays in MDCK cells in plaque forming units per mL (PFU/mL) including the inoculum, labelled Inoc. Each point represents the mean of three biological replicates (n=3). Statistical significance was calculated in comparison to the wildtype avian virus (referred to as BC HA 190E 226Q) at each time point in GraphPad Prism 10.0.3 using log-transformed titres in a mixed effects model, with the Sidak method for multiple comparisons with titres below the limit of detection (50 PFU/mL) transformed as half the limit of detection. In MDCKs **B**. BC HA 190E 226Q had statistically significantly higher growth than all other viruses at 16h.p.i. and 24h.p.i., and BC HA 190E 226H and BC HA 190D 226H at 48 and 72h.p.i. In Calu-3 cells **C**. BC HA190E 226Q growth was statistically significantly higher than BC HA 190E 226H at 48h.p.i, and BC HA 190D 226H at 24, 48, and 72h.p.i. Statistical significance is reported as p-values (**Supplemental Table 1**) represented by an asterisk, where p ≤ 0.05 (*), p ≤ 0.01 (**), p ≤ 0.001 (***), p ≤ 0.0001 (****). **D.** In HNECs, BC HA 190E 226Q had statistically significant growth over the double mutant BC HA 190D 226H at 48 and 72h.p.i and over the single mutant BC HA 190E 226H at 72h.p.i.

Viruses were first used to infect MDCK cells at a low multiplicity of infection (MOI) of 0.01 to quantify multicycle replication kinetics (**Figure 1B**). The wildtype avian virus, BC HA 190E 226Q, reached significantly higher titres than all mutant viruses at 16 and 24 hours post infection (h.p.i.) (**Supplemental Table 1a**). At later time points, BC HA 190D 226Q achieved higher titres than BC HA 190E 226H and the double mutant, reaching equivalent titres to the wildtype by 72h.p.i.

Next viral replication kinetics were measured in immortalized human lung cells (Calu-3). Again, the wildtype virus BC HA 190E 226Q had a replicative fitness advantage in these cells over BC HA 190E 226H and BC HA 190D 226H (**Figure 1C, Supplemental Table 1b**). However, the BC HA 190D 226Q mutant had equivalent replication kinetics to the wildtype BC HA 190E 226Q in Calu-3 cells.

The replication kinetics of these four recombinant viruses was then tested in highly differentiated human nasal epithelial cells cultured with an air-liquid interface and containing a mixture of ciliated and non-ciliated cells, resulting in a layer of secreted mucus at the apical surface. These cultures serve as an in vitro model to measure virus fitness in the environment of the human airway. In these cultures, the double mutant BC HA 190D 226H replicated only poorly, achieving greatly reduced titres by 48 and 72h.p.i (**Figure 1D**). On the other hand, replication of either of the single mutants was less impacted and titres were similar to wildtype except at 72h.p.i., when the BC HA 190E 226H titre was slightly lower.

### Mutant viruses exhibit weaker binding of sialic acid than wildtype virus

We then used the four recombinant viruses to investigate if these mutations influenced the sialic acid preference and binding affinity of HA. The receptor binding properties of these viruses were first investigated using purified viruses and sialoglycan receptor analogues via biolayer interferometry (BLI). α2,3-sialyllactosamine (3’SLN) was used as a typical avian receptor analogue and α2,6-sialyllactosamine (6’SLN) to represent the human receptor analogue. The wildtype avian virus, BC HA 190E 226Q retained binding to the avian analogue 3’SLN, while the human virus control, A/England/195(H1N1) showed only detectable binding to the human analogue 6’SLN (**Figure 2A**). BC HA 190E 226H had reduced binding to 3’SLN, with no detectable binding to 6’SLN. No binding could be detected by BLI for BC HA 190D 226Q nor for BC HA 190D 226H to either receptor analogue.

**Figure 2.**
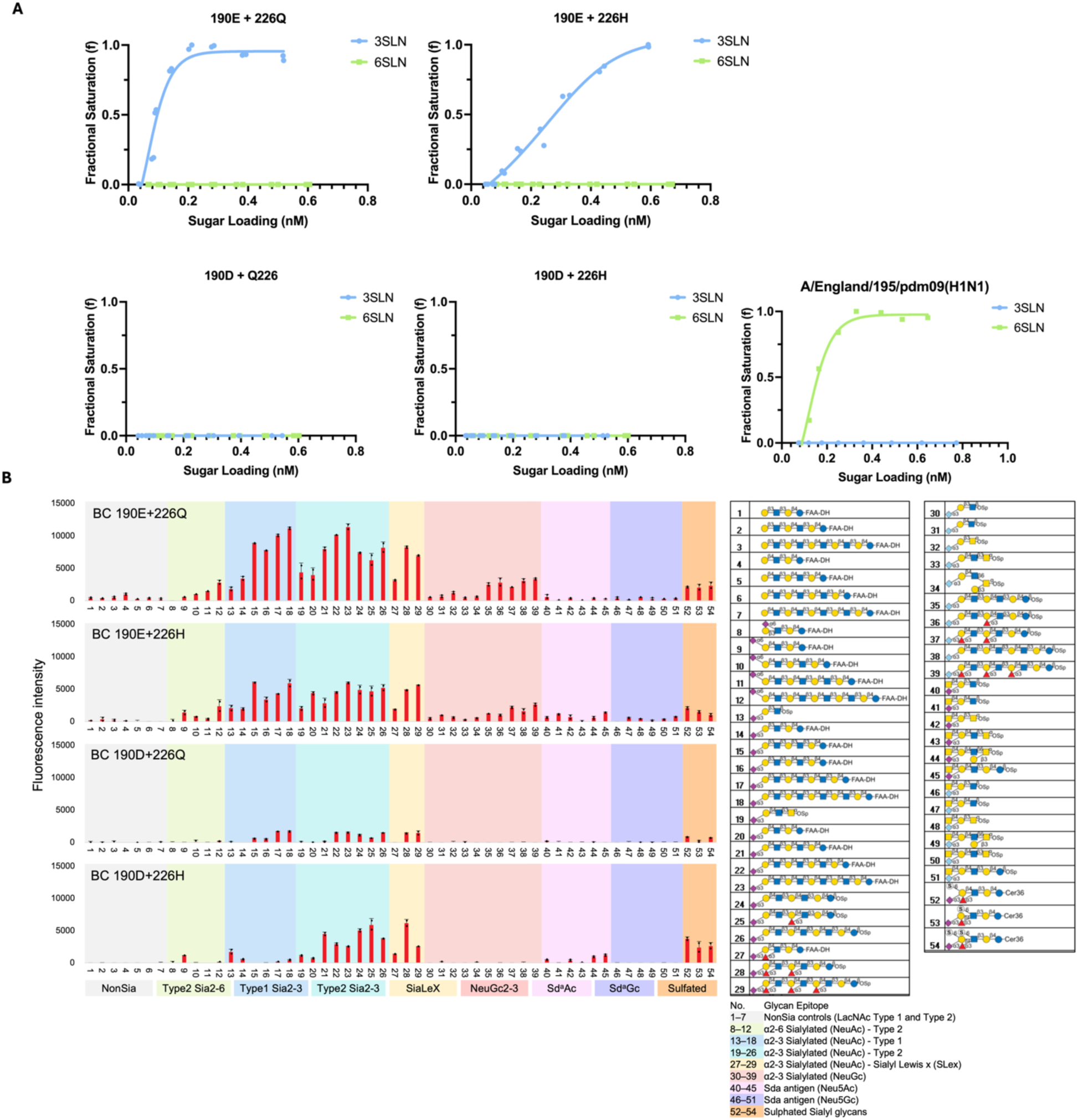
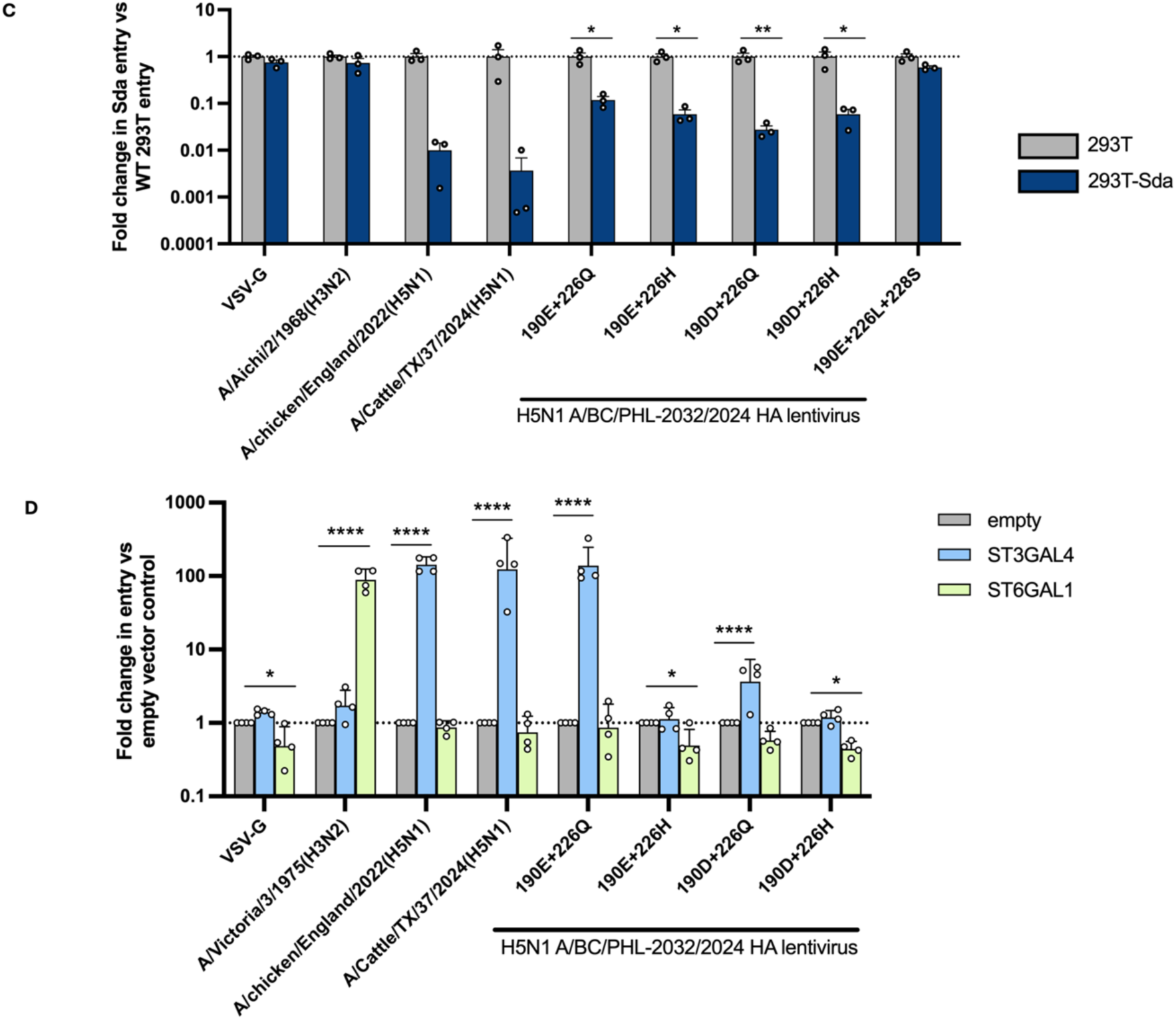
Mutant viruses exhibit reduced sialic acid affinity to α2,3-linked sialic acids. **A.** The receptor binding of four viruses BC HA 190E 226Q, 190E 226H, 190D 226Q, 190D 226H measured by BLI using receptor analogues 3’SLN (avian) and 6SLN (human). A/England/195/pdm09(H1N1) is a control which binds only 6SLN. **B.** Glycan microarray binding profiles of the four BC virus samples. Data represent the mean fluorescence intensities of duplicate glycan probe spots printed at 5 fmol per spot. Error bars indicate half the difference between the duplicate measurements. The 54 lipid-linked glycan probes are grouped into non-sialylated and sialylated glycans according to their structural features, as indicated by the colour panels. The complete list of glycan probes and binding scores are provided in **Supplementary Data File 1**. **C.** Pseudovirus entry assays into wildtype and SDA-expressing 293T cells with HA-expressing pseudotyped lentiviruses made with exogenous bacterial NA (**Supplemental Figure 2**). Cell entry was plotted as fold change in entry into 293T-Sda expressing cells divided by entry into wildtype 293T cells. **D.** Pseudovirus entry in sialyltransferase knockout 293 cells (ΔST3GAL1/2/3/4/5/6 ΔST61/2) transiently transfected with ST3GAL4 or ST6GAL1, which allow for the expression of α2,3-linked SA and α2,6-linked SA expression, respectively^29,30^. Results shown are the mean of at least three independent biological repeats (in duplicate). VSV-G pseudovirus and (**C**) A/Aichi/2/1968(H3N2) or (**D**) A/Victoria/3/1975(H3N2) are controls which do not enter via α2,3-linked SA and enter each cell line equally. A/chicken/England/2022(H5N1) and A/Cattle/TX/37/2024(H5N1) are controls which are expected to use α2,3-linked SA for entry. BC HA pseudovirus with the mutations Q226L and G228S was generated as a positive control for an H5 HA that would use α2,6-linked SA^21^. All other BC mutants had a statistically significant reduction entry into the SDA-expressing cells using (**C**) multiple t-tests or (**D**) two-way ANOVA in GraphPad Prism with Šídák’s multiple comparisons test. Statistical significance is reported as p-values (**Supplemental Table 2**) represented by an asterisk, where p ≤ 0.05 (*), p ≤ 0.01 (**), p ≤ 0.001 (***), p ≤ 0.0001(****).

The panel of purified recombinant viruses were then tested using a sialoglycan-focused microarray containing 54 lipid-linked glycan probes (**Figure 2B**). This array includes non-sialylated LacNAc control glycans of both Type 1 (Galβ1-3GlcNAc) and Type 2 (Galβ1-4GlcNAc) backbone structures, as well as a diverse set of sialylated glycans featuring both α2,6- and α2,3-linked Neu5Ac structures on Type 1 and Type 2 backbones. It also incorporates structurally distinct motifs such as sialyl Lewis x (SLex), Neu5Gc-containing glycans, and Sda antigens (Neu5Ac and Neu5Gc forms). In addition, three sulfated sialyl glycans are included to capture charge-dependent interactions. As in the BLI, BC HA 190E 226Q and BC HA 190E 226H primarily bound α2,3-linked sialoglycans, with the single mutant BC HA 190E 226H exhibiting reduced binding compared to 190E 226Q. BC HA 190D 226Q and BC HA 190D 226H showed markedly reduced sialic acid binding overall compared to the wildtype. This decreased binding to α2,3-linked sialoglycans for the single and double mutant viruses was even more evident when using recombinant HA protein, likely because of a loss of avidity effects of a multivalent interaction when the full viruses were used (**Supplemental Figure 1**).

To study the entry phenotype caused by the HA mutations, we generated HA-expressing pseudotyped lentiviruses^26,27^. These included the previous four combinations of HA mutations: 190E 226Q, 190E 226H, 190D 226Q, 190D 226H, and a further mutant, 190E Q226L G228S. The new mutant was used to indicate the expected properties for an H5 HA that has undergone human host adaptation, since previous studies have shown a receptor switch to α2,6-SA of H5 HA with the Q226L mutation alone or in combination with G228S^21,22^. Introducing these mutations into a single cycle replication-deficient system, such as a pseudovirus means that they can be studied safely in the laboratory. These pseudoviruses were tested for cell entry in wildtype human embryonic kidney (HEK) 293T cells and HEK 293T cells stably expressing the B4GALNT2 enzyme, which results in the blocking of α2,3-linked SA containing glycans to create the Sda epitope thereby inhibiting entry of α2,3-dependent HAs **(Figure 2C, Supplemental Figure 2**)^28^. pseudovirus bearing the mutant BC HA 190E Q226L G228S, VSV-G (a non-sialic acid dependent viral glycoprotein), or a human H3N2 (A/Aichi/2/1968) were not significantly inhibited in their entry into the Sda-expressing cells. This finding is consistent with these pseudoviruses not being dependent on α2,3-linked SA for entry. Entry of all other BC HA-pseudoviruses [190E 226Q, 190E 226H, 190D 226Q, 190D 226H] was statistically significantly reduced in the Sda-expressing cells compared to entry into wildtype 293T cells. This data aligns with microarray data to suggest viruses with these HAs can use α2,3-linked SA as receptors, even if their affinity for sialic acid in general has been reduced.

We then used a similar assay in which 293 cells knocked out for sialyl-transferase expression were reconstituted with either α2,3-SA or α2,6-SA by re-expressing sialyl-transferase enzymes^29^. In this assay, a similar pattern was seen whereby the mutations E190D and Q226H in HA alone or in combination did not enable α2,6-SA mediated cell entry (**Figure 2d**). Although both mutations individually, or in combination, resulted in a significant decrease in α2,3-SA-dependent entry relative to the wildtype virus. sialyltransferase knockout 293 cells (ΔST3GAL1/2/3/4/5/6 ΔST61/2)

### E190D and Q226H provide a fitness advantage in escaping inhibition by human mucus

Thus far, the emerging HA mutations detected in clinical samples did not confer human-like receptor binding properties, instead only decreasing binding to avian-like α2,3-SA receptors. To explore what evolutionary pressures might be driving such a change, we tested mutant HA interaction with human airway mucus, which is known to exert a protective anti-viral effect against influenza viruses^31^. Human airway mucus was obtained through washing the apical surface of the primary cultures of human nasal epithelial cells and was then mixed with a standardized titre of each virus. Remaining infectivity was determined by plaque assays on MDCK cells (**Figure 3A**).

**Figure 3.**
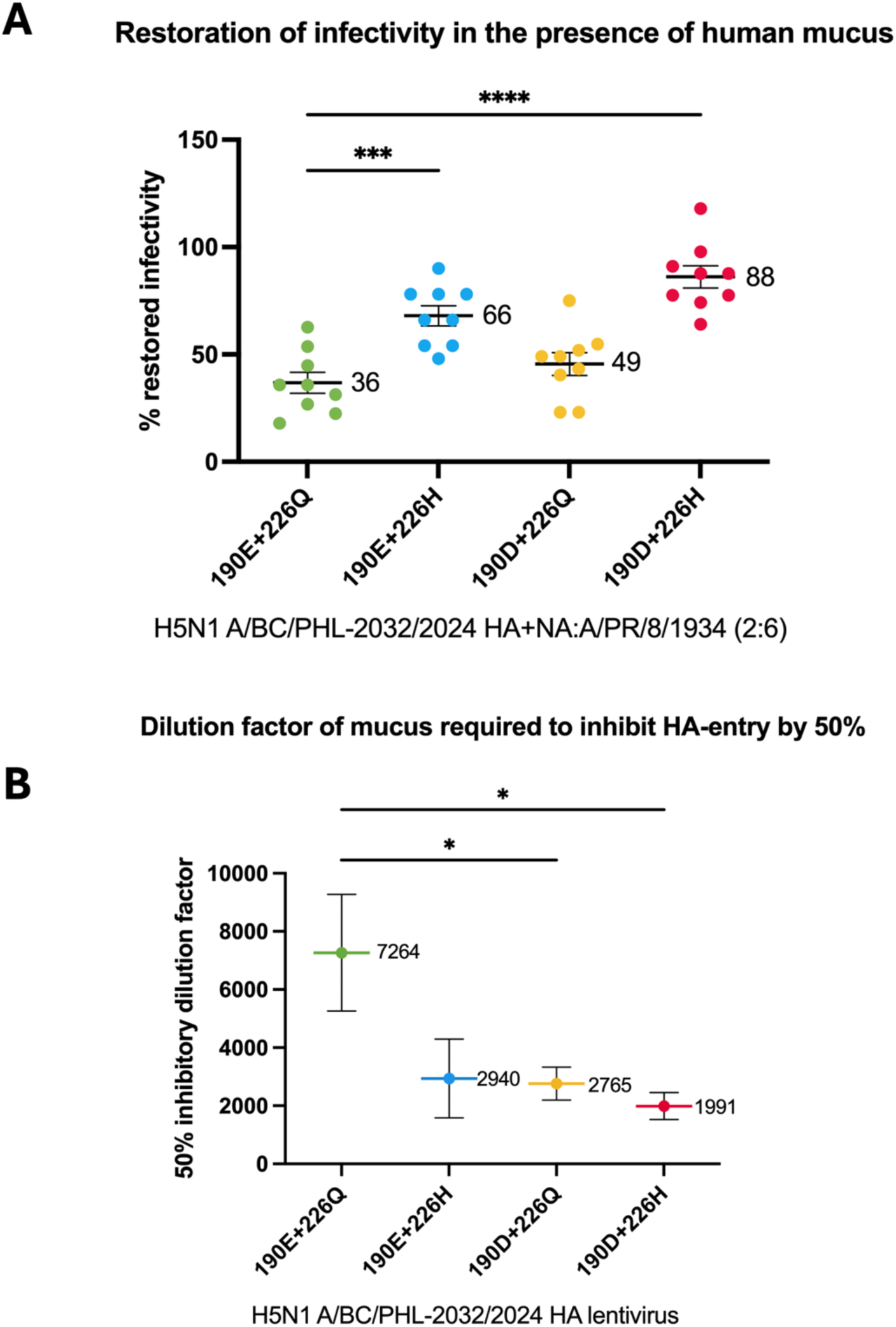
E190D and Q226H provide a fitness advantage for escape from human mucus. **A.** H5N1 A/BC/PHL-2032/2024 HA+NA: H1N1 A/Puerto Rico/8/1934(2:6) viruses were incubated with human mucus diluted 1:70 in PBS. Mucus was diluted 1:70 in PBS and incubated with virus. Mucus and virus containing tubes were incubated on ice for 2hrs (n=9). Mean percent restored infectivity is labelled on the graph with mean and standard deviation shown. Mixture was added to MDCK cells and incubated at 33°C for 2hrs before being removed. Virus growth in the presence of mucus was normalized to a no-mucus control which was used to calculate restoration of infectivity. In these assays, BC HA 190E 226Q had a reduced ability to restore infectivity over BC HA 190E 226H and BC HA 190D 226H. Mean percent restored infectivity is labelled on the graph for each virus with mean and standard deviation shown (n=9). **B.** Mucus neutralisation assays were performed with HA-expressing pseudoviruses with the four BC HA mutants. BC HA 190E 226Q required the lowest concentration of mucus to reduce pseudovirus cell-entry by 50%, which was statistically significant when compared to BC HA 190D 226Q and 190D 226H. Statistical significance between groups was compared in GraphPad Prism via a one-way ANOVA with Dunnett’s multiple comparisons test. An asterisk indicates statistical significance where p ≤ 0.05 (*), p ≤ 0.01 (**), p ≤ 0.001 (***), p ≤ 0.0001 (****) (**Supplemental Table 3**).

Human airway mucus can inhibit the infectivity of influenza viruses by competitively inhibiting the HA-SA receptor interaction on target cells. The NA on virus particles can cleave soluble decoy receptors in mucus and restore virus infectivity at physiologically relevant temperatures. In these assays, all three mutant viruses were less inhibited by the mucus than the avian-like wildtype BC HA 190E 226Q, significantly so for BC HA 190E 226H and for BC HA 190D 226H (**Figure 3A**).

The human airway mucus was also titrated for its ability to inhibit the entry of BC HA-pseudotyped lentiviruses. In this assay, we generated a mucus neutralisation curve to establish any quantitative difference in neutralization effect. Mucus was serially diluted (as human serum would be in a traditional neutralisation assay), and the dilution of mucus that inhibited 50% pseudovirus entry was calculated and reported as an IC50 value (**Figure 3B**). Pseudovirus bearing the BC HA 190D 226H was the least sensitive to mucus, with around four times more mucus needed to reach the 50% inhibitory concentration than BC HA 190E 226Q. As with the mucus inhibition assays with the recombinant influenza viruses (**Figure 3A**), the highest concentration of mucus was needed to reduce BC HA 190D 226H cell entry by 50%, suggesting the effect of E190D and Q226H was additive in these assays.

### Mutant viruses are less sensitive to oseltamivir and zanamivir than wildtype

It was reported that both patients in British Columbia and Louisiana were treated with oseltamivir upon admission to hospital^18,19^. According to the initial case report, the 190D and 226H minor allele frequency mutations in the British Columbia case were identified in sequences taken after oseltamivir treatment began^18^. Therefore, viral replication kinetics studies were repeated in the presence of oseltamivir, as previous studies have indicated that HA mutations which decrease sialic acid binding affinity could facilitate oseltamivir escape^32^.

In the presence of oseltamivir and zanamivir, the fitness advantage for replication kinetics shifted away from the wildtype virus in all cell lines. Titres of the wildtype virus were significantly reduced in all cell lines at the majority of timepoints in the drug treated conditions as compared to the untreated condition (**Figure 4**). The mutant viruses remained unaffected by treatment with oseltamivir or zanamivir across most timepoints and cell lines. When comparing wildtype virus replication side by side with mutant virus replication under drug treatment, the wildtype had reduced viral titres (**Supplemental Figure 3**).

**Figure 4.**
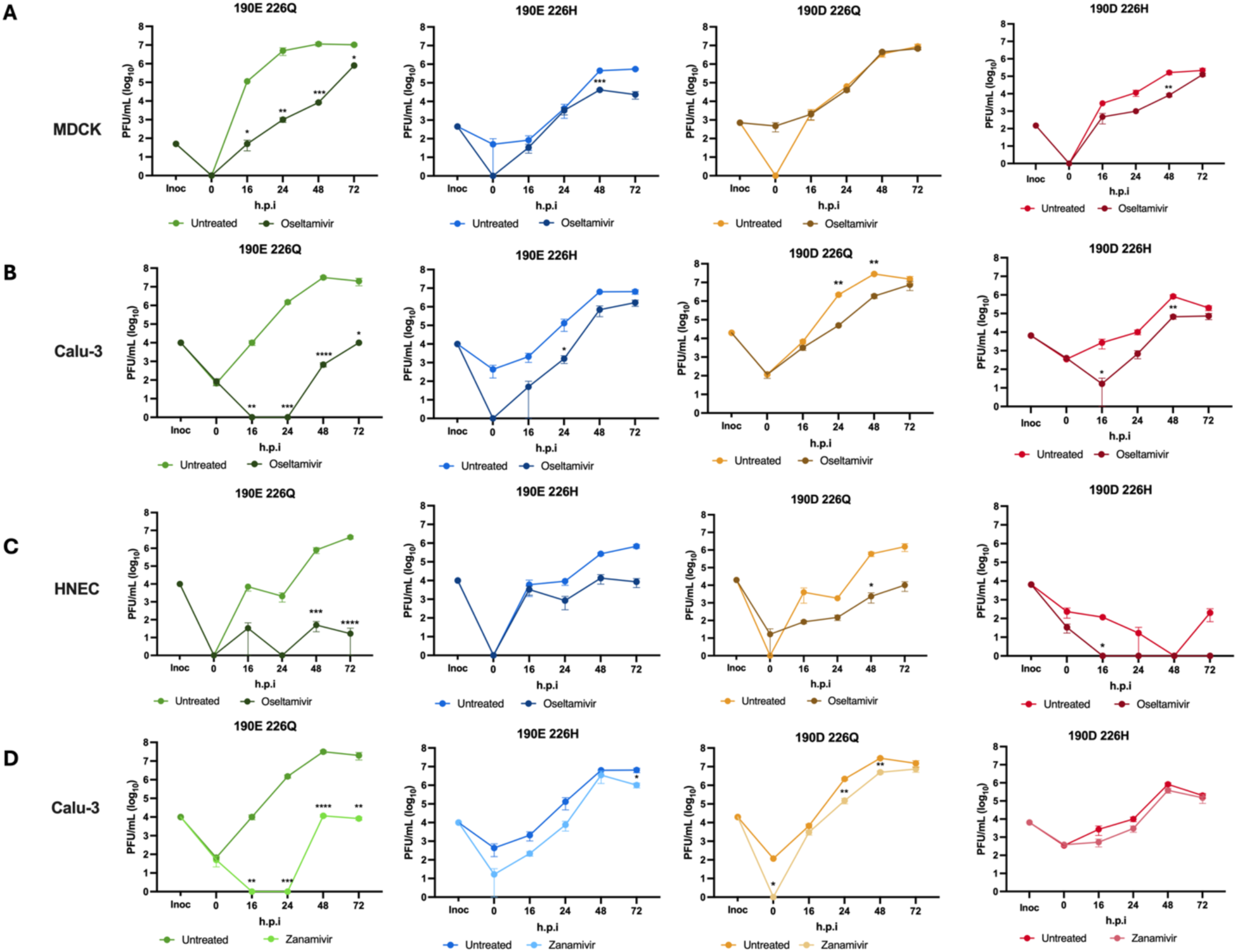
Fitness advantage shifts towards mutant viruses in the presence of oseltamivir and zanamivir. Viral growth kinetics of the H5N1 A/BC/PHL-2032/2024 HA+NA: H1N1 A/Puerto Rico/8/1934(2:6) viruses. Supernatant was collected from cells at 0, 16, 24, 48, and 72h.p.i. Viral titres were quantified by plaque assays in MDCK cells in PFU/mL including the inoculum, labelled Inoc using viruses at MOI 0.01. Each point represents the mean with of three biological replicates (n=3) with the standard deviation. Untreated conditions have been replotted from Figure 1 to compare with treated conditions, which were run on the same day. **A.** In MDCK cells, at 16, 24, 48, and 72h.p.i. BC HA 190E 226Q had statistically significantly reduced titres in the presence of oseltamivir. The mutant viruses had the same titres with and without oseltamivir treatment, except at 48h.p.i for 190E 226H and 190D 226H. **B.** In Calu-3 cells, BC HA 190E 226Q had statistically significantly reduced titres in the presence of oseltamivir at 16, 24, 48, and 72h.p.i. The untreated condition had higher tires at 24h.p.i. for 190E 226H, 24 and 48h.p.i. for 190D 226Q, and 16 and 48h.p.i. for 190D 226H. **C.** In HNECs, the wildtype had reduced titres in the presence of oseltamivir at 48 and 72h.p.i. 190D 226Q had reduced titres at 48h.p.i. and 190D 226H had reduced titres at 16h.p.i. **D.** In the presence of Zanamivir in Calu-3 cells, 190E 226Q had reduced tires at 16, 24, 48, and 72h.p.i. 190E 226H had reduced titres at 72h.p.i, and 190D 226Q had reduced titres at 0, 24, and 48h.p.i. Statistical significance for viral replicative fitness was calculated as in Figure 1. Statistical significance is reported as p-values (**Supplemental Table 4**) represented by an asterisk, where p ≤ 0.05 (*), p ≤ 0.01 (**), p ≤ 0.001 (***), p ≤ 0.001 (****).

## Discussion

In this study, we sought to take a comprehensive approach to analysing the possible evolutionary advantages for the appearance of minor frequency alleles in human H5N1 genotype D1.1 cases: E190D and Q226H. While others have reported that these viruses do not have a fitness advantage in regards to growth^33^ (without antivirals present), the independent appearance of these mutations in two unlinked cases suggests that there must have been an advantage for the mutations to emerge, particularly at two sites historically associated with host adaptation^20–22^.

When we investigated the sialic acid binding affinity of these mutants using multiple complementary approaches, we found that both 190D and 226H reduced binding to sialylated glycans, in line with several other reports^33–35^, but in contrast to a recent preprint which suggests that Q226H could bind α2,6-linked SA^36^. Our glycan microarray analysis also suggested that a very low level of α2,3-linked SA binding affinity is retained.

IAV must escape sialic acid decoy receptors in the respiratory tract to prevent viral sequestration by mucus^37–40^. Mucus escape or sensitivity is typically considered to be a role of NA^41^; however, the results here suggest that HA can also affect mucus escape. When neutralisation by mucus was tested with all four variants, the wildtype virus was the most sensitive. The double mutant was the least sensitive, and the individual mutations 190D and 226H had additive effects in both pseudovirus and fully replicating viral systems. These changes in mucus sensitivity are mapped specifically to HA, as that was the only change between the viruses tested, suggesting that HA may have a previously uncharacterized role in determining mucus susceptibility and escape from decoy receptors.

While the HA mutations 190D and 226H may have emerged to promote escape from human mucus, we hypothesised that this mechanism was also driven by treatment with oseltamivir. Indeed, in the presence of oseltamivir or zanamivir, mutant viruses replicated to higher titres in all cell lines. This suggests that the lack of sensitivity to oseltamivir and zanamivir is HA-mediated in BC H5N1 viruses.

Although oseltamivir is a neuraminidase inhibitor, there is precedent for HA-mediated oseltamivir escape. Zhang et al. identified a single HA mutation (HA-K130N or K130E) in pdm09 H1N1 viruses which have been circulating since 2019 that reduced oseltamivir sensitivity^32^. One possible explanation for the emergence of the HA mutations could be that continued exposure to an NA inhibitor resulted in reduced effective D1.1 NA activity, and that the HA 190D and 226H mutations developed to compensate for this effect to maintain effective HA/NA balance. Lower NA activity and reduced sialic acid affinity could serve in some cases as a previously misunderstood advantage to escape antivirals and host decoy sialic acid receptors in human mucus.

In this study, we have highlighted the understudied role of HA in two functions which have often been associated with NA: antiviral escape and a lack of sensitivity to human mucus. This suggests that co-evolution between the functional roles of HA and NA should be considered in tandem with emerging strains to greater understand and monitor the possibility of oseltamivir-resistant viruses that can overcome host barriers, such as mucus. Further work should seek to monitor not only well-known oseltamivir resistance mutations in NA but should also characterize HA mutations as drivers of oseltamivir resistance and consider the inverse relationship between NA activity and HA-driven mucus escape.

## Methods

### Ethical Approval

Experimental procedures involving the use of embryonated chicken eggs for the propagation of viruses and polysera preparation in chickens were undertaken in accordance with UK Home Office regulations (license numbers PP671846 and PP0142098). Approval to perform these procedures was approved by the animal welfare ethical review boards at The Pirbright Institute and Imperial College London.

### Viruses

The HA and NA sequences of A/British Columbia/PHL-2032/2024(H5N1) (referred to as H5 BC or BC HA) were acquired from Jassem et al.^17,18^ via GISAID (EPI_ISL_19548836)^42,43^, synthesized by GeneScript, and cloned into a pHW2000 vector. The multi-basic cleavage site of the HA was replaced by a monobasic site, as previously described^44^. Virus rescues were performed using the remaining gene segments, PA, PB1, PB2, NP, M, and NS, from the laboratory-adapted strain A/Puerto Rico/8/1934(H1N1) (referred to as PR8). BC HA viruses were propagated in MDCKs or 10-day old embryonated chicken eggs to achieve a higher titre. Sequences were confirmed before use in assays (Full Circle Biolabs).

### Site directed mutagenesis (SDM)

Mutations in HA-containing plasmids were generated using site directed mutagenesis. For HA plasmids used in virus rescues, the single-site mutagenesis QuikChange II kit (Aligent #200523) was used. HA plasmids used to generate pseudovirus were mutated with the Q5 Site-Directed Mutagenesis kit (NEB #E0554S) and confirmed via whole plasmid sequencing (Full Circle Labs). Both kits were used according to the respective manufacturer’s instructions.

### Plaque assay

Infectious virus was titrated by plaque assays on MDCK cells, as described previously^45^. Viruses were serially diluted 10-fold and used to inoculate MDCK cells. Cells were overlaid with 2% agarose in with 2µg/mL N-tosyl-L-phenylalanine chloromethyl ketone (TPCK) trypsin (Sigma) in DMEM supplemented with 1% NEAA, 3% Penicillin/Streptomycin, 20% 10X MEM, 5.6% BSA frac. V, 2% 1mM L-Glutamine, 4% Sodium Bicarbonate, 2% HEPES (all Gibco), 1% DEAE (Sigma), and 103.4% sterile H_2_O at one hour post infection (h.p.i.). Plaques were developed after incubation at 37°C for 72hrs using crystal violet stain.

### Viral replication kinetics in immortalized cell lines

Viral replication kinetics were measured in MDCK and Calu-3 cells, seeded 1:4 in a 12-well plate. Cells were inoculated with a multiplicity of infection (MOI) of 0.01 calculated based on plaque assay results. At one hour post infection, virus was replaced with serum free media and incubated at 37°C. Culture supernatants containing virus were collected at 0, 16, 24, 48, 72 hours post infection (h.p.i) and titrated by plaque assays in MDCK cells as above. The growth data plotted is the mean of three independent experiments. Viral growth curves were repeated in the same manner in the presence of 10uM oseltamivir or 10μM zanamivir (Calu-3 cells only).

Statistical significance was calculated in comparison to the wildtype avian virus (referred to as BC HA 190E 226Q) at each time point in GraphPad Prism 10.0.3 using log-transformed titres in a mixed effects model, using the Sidak method for multiple comparisons with titres below the limit of detection (50 PFU/mL) transformed as half the limit of detection. Statistical significance is reported as p-values (**Supplemental Table 1, 5**).

### Viral replication kinetics in primary human nasal epithelial cell cultures

Viral replication kinetics were measured in human nasal epithelial cultures (HNECs) from a pool of 14 healthy human donors (Epithelix MucilAir™-Pool nasal, batch: MP0012). Upon receipt of HNECs, supplier instructions were followed to remove cells from agarose. HNECs were cultured in 24-well plates. Cells were washed with PBS and given fresh media every 48hrs, using MucilAir™ medium (Epithelix). Mucus was collected in PBS once per week and used for mucus assays.

Cells were inoculated with a MOI of 0.01 calculated based on plaque assay results. At one hour post infection, virus was replaced with serum free media and incubated at 37°C. Culture supernatants containing virus from MOI 0.01 conditions were collected at 0, 16, 24, 48, and 72h.p.i and titrated by plaque assays in MDCK cells. The growth data plotted is the mean of three independent experiments. Viral growth curves were repeated in the same manner in the presence of 10uM oseltamivir, which was added to the basal side of the transwell.

As above, statistical significance was calculated in comparison to the wildtype avian virus (referred to as BC HA 190E 226Q) at each time point in GraphPad Prism 10.0.3 using log-transformed titres in a mixed effects model, using the Sidak method for multiple comparisons with titres below the limit of detection (50 PFU/mL) transformed as half the limit of detection. Statistical significance is reported as p-values (**Supplemental Table 1, 5**).

### Bio-layer interferometry (BLI)

After propagation, viruses were purified through 30% and 60% sucrose gradients at 27,000rpm for 2hrs at 4°C. Purified viruses were normalised by viral nucleoprotein (NP) enzyme-linked immunosorbent assays (ELISA) to 100pmol^46^. Viruses were diluted with 10μM oseltamivir carboxylate (Roche) and 10μM zanamivir (GSK) against avian-like sugar analogue (3SLN) and human-like sugar analogue (6SLN). Binding affinity of each virus to the sugar analogues was tested using streptavidin biosensors with the Octet® R8 system (Sartorius). As previously described, virus binding to each sugar analogue was normalised to fractional saturation the concentrations of sugar loading^47^.

### Antisera preparation

Chicken polyclonal antiserum against the BC 190E 226Q virus was generated in embryonated chicken eggs. The harvested allantoic fluid was inactivated with 0.1% (v/v) β-propiolactone (BPL), and complete inactivation was confirmed by passaging the antigen three times in embryonated chicken eggs, followed by haemagglutination (HA) assay. The inactivated virus was then concentrated by ultracentrifugation at 27,000 rpm for 2 h. Three-day-old specific-pathogen-free (SPF) chickens were inoculated with 1,024 haemagglutinating units (HAU) of the concentrated inactivated virus emulsified with the oil emulsion adjuvant Montanide (Seppic) at an adjuvant-to-virus ratio of 7:3. A booster dose was administered 10 days after the primary inoculation.

### Glycan Microarrays

The binding specificity of the virus and HA samples was assessed using a focused neoglycolipid (NGL)-based, sialyl glycan array set comprising 54 lipid-linked oligosaccharide probes. The full list of probes and their sequences is provided in **Supplementary Data File 1**.

Microarray analyses of live influenza viruses were performed essentially as described (Liu, et al., 2010)^48^. In brief, after blocking the array slides with 2% (w/v) bovine serum albumin (BSA, Sigma A7030) in Hepes buffered saline (HBS; 5 mM Hepes, pH 7.4, 150 mM NaCl, 5 mM CaCl_2_), live virus suspension (100pmol quantified by NP ELISA as described above^41^) was incubated for 90 minutes at room temperature in the presence of Zanamivir and Oseltamivir (100 µM each). Following fixation with 4% paraformaldehyde, bound virus was detected using the chicken polyserum raised against BC E190 Q226 virus (1:200) followed by biotinylated anti-chicken IgY (1:200; Abcam, ab207998) for 60 minutes and Alexa Fluor 647-conjugated streptavidin (Molecular Probes) at 1 µg/mL. The analyses of HA proteins were essentially as described in Liu, et al., 2012^49^. Microarray slides were blocked with 1% BSA (w/v) in HBS. Recombinant C-tagged HA was precomplexed with biotinylated anti-C-tag antibody (Thermo Scientific 7103252500) in a ratio of 1:1 and overlaid onto arrays at 40 µg/mL for 90 min incubation at room temperature. Binding was detected with Alexa Fluor 647-conjugated streptavidin (1 µg/mL). As diluent the blocker solution was used.

Additional details regarding the glycan library, microarray fabrication, imaging procedures, and data analysis are provided in the Supplementary Glycan Microarray document (**Supplementary Table 6**), in accordance with the Minimum Information Required for A Glycomics Experiment (MIRAGE) guidelines for reporting glycan microarray data^50^.

### Production of cell lines

Pseudotyped lentiviruses were produced containing a B4GALNT2 gene as follows. HEK 293T cells were transfected in a six-well plate with lipofectamine 3000 (Invitrogen), 0.4µg of HIV packaging plasmid (pGAG-POL), 0.4µg VSV-G, and 0.4µg B4GALNT2 plasmids. B4GALNT2 catalyses the last step of synthesis of the Sda epitope, which modifies α2,3-linked SA to become inaccessible to bind HA (kindly provided by Professor Nicholas S. Heaton, Duke University)^51^. Supernatant with pseudoviruses was collected 48hrs post transfection and filtered through a sterile 0.45-µm Millex®-HA filter unit (Millipore, SLHA033SB).

B4GALNT2 containing pseudoviruses were diluted 1:10 in DMEM with 10% FBS, 1% pen-strep, 1% NEAA and added to a six-well plate seeded with wildtype 293T cells for B4GALNT2. These cells will be referred to as 293T-Sda.

Cells were treated with 4µg/ml of puromycin (selection antibiotic) at 48hrs and 96hrs post transfection. Cells were then treated with 0.05% Trypsin-EDTA (1X) (25300-054) and seeded into 25cm^2^ cell culture flask (Grenier, 690175) with 1µg/ml of puromycin.

### Immunofluorescence of cell lines

The 293T-Sda cells were validated by immunofluorescence. Cells were stained as described previously^52^. In brief, cells were seeded in a twelve well plate. After 24hrs, media was removed from the cells. Cells were fixed with 200μL 4% paraformaldehyde in PBS for 30min, and permeabilized for 10min with 200μL 1% Triton X-100 in PBS (Sigma #X100). Cells were washed with 1X PBS and stained depending on the cell type.

293T-Sda cells were stained with 20ug/mL lectins conjugated to fluorescein (FITC) in 1X OptiMEM, either MAL I (Vector Laboratories, FL-1311-2), SNA (Vector Laboratories, FL-1301-2), or Dolichos biflorus lectin (DBA; binds the Sda epitope, Sigma-Aldrich). Cells were incubated with the lectins for 1hr at 4°C, then washed once with 200μl OptiMEM. DBA lectin was conjugated to biotin, so after 1hr these wells were washed and stained with streptavidin-conjugated Alexa Fluor™ 488nm (Invitrogen, S11223) for 1hr at 4°C.

Cells were stored in PBS and visualised with an EVOS M3000 Imaging Auto fluorescent microscope. The 293T-Sda cells showed reduced MAL-I staining compared to the wildtype cells, suggesting that Sda was successfully blocking a2,3-linked sialic acids (**Supplemental Figure 2a**).

### Flow cytometry analysis of cell lines

Cell surface expression for 293T-Sda cells was also quantified by flow cytometry. Cells were stained with Zombie-NIR fixable viability dye (BioLegend, 423105) to determine live vs dead populations. Cells were then fixed as in the immunofluorescence studies above. Cells were stained with lectins as in immunofluorescence above.

Events were then acquired on the NovoCyte Penteon flow cytometer (Aligent Technologies). A minimum of 10,000 events was collected per condition, and results were gated to exclude doublets and dead cells (**Supplemental Figure 2b**). As with immunofluorescence, the Sda cells had reduced MAL-I staining with unaffected SNA staining, supporting the idea that α2,3-linked sialic acids were successfully modified by Sda (**Supplemental Figure 2c**).

### Production of pseudoviruses

HA-expressing pseudotyped lentiviruses were produced in HEK293T cells seeded in 6-well plates as previously described^26,27,30^. Cells were transfected with lipofectamine 3000 (Invitrogen), 0.4µg HA or VSV-G, 0.4µg of the HIV packaging plasmid expressing *gag* and *pol*^53^, 0.6µg of a luciferase reporter construct, and 0.2 µg of TMPRSS11D (H3 only). The following plasmids were generated in pcDNA3.1(+) by GenScript: H3 HA – A/Aichi/2/1968 (H3N2; BAF37221.1), and human TMPRSS11D/HAT (EAX05559.1).

A/British Columbia/PHL-2032/2024(H5N1)^17,18^ (EPI_ISL_19548836 obtained from GISAID^42,43^) was generated in pcDNA3.1(+) by Twist Bioscience.

After 24hrs, one unit of neuraminidase from *Clostridium perfringens* (Sigma-Aldrich, N2876) was added with fresh media to induce virus budding. Pseudovirus containing supernatant was collected after 48hrs and filtered through a sterile 0.45-µm Millex®-HA filter unit. All pseudoviruses were titrated before use in entry assays as previously described^27^. For oseltamivir sensitivity assays, pseudoviruses were produced in the same manner as above with 0.2 µg H5N1 NA co-transfected with HA protein instead of bacterial neuraminidase.

### Pseudovirus entry assays

Pseudoviruses were added in duplicate to a white Nunc™ MicroWell™ 96-well, flat-bottom microplate with 50µL pseudotyped influenza virus per well. Twofold serial dilutions of pseudoviruses in media were used to test pseudovirus entry into cell lines. The starting dilution was based on the virus titrations. Viruses were diluted to reach a standard titre of 1×10^6^ RLU after 48hrs. All entry assays were run with three independent biological repeats. Pseudoviruses were incubated at 37°C with 50µL of 1×10^5^ cells. Cell lines tested for pseudovirus entry included wildtype HEK239T cells and HEK293T-SDA cells.

Each plate included a cell-only control. VSV-G was used as a control for sialic acid independent entry, A/Aichi/2/1968(H3N2) was used as a positive control for α2,6-linked sialic acid entry. 48hrs post-transduction cells were lysed with Bright-Glo^TM^ Luciferase Assay System (Promega). The RLU of the cell lysate was determined using the FLUOstar OMEGA microplate reader (BMG Labtech).

Sialyl-transferase reconstitution assays were adapted as previously described^30^. 293 cells lacking endogenous sialyl-transferase expression (HEK293 KO: ΔST3GAL1/2/3/4/5/6, ΔST61/2^29^) were transiently transfected with 100 ng of expression plasmids encoding empty vector, human ST3GAL4, or ST6GAL1, and mixed with 400 ng of empty vector. 24hrs later cells were harvested, diluted to 105 cells/ml, transferred into 96 well plates (100μL/ well) and mixed with 100μL of pseudovirus. Cells were then left for 48hrs at 37°C, 5% CO_2_ then lysed and read using Bright-Glo^TM^ Luciferase Assay System and a Glomax discover plate reader (Promega). Data for each pseudovirus was normalised to the empty vector control.

Differences between entry into cell lines were plotted as fold change relative to the wildtype cells. Means of each condition were compared per pseudovirus using a two-way analysis of variance (ANOVA) test. An asterisk indicates statistical significance where *p* ≤ 0.10 (.), *p* ≤ 0.05 (*), *p* ≤ 0.01 (**), *p* ≤ 0.001 (***).

### Mucus inhibition assays

Mucus was collected once per week from HNECs in 200µL PBS per transwell. Mucus was diluted 1:70 in PBS and incubated with virus. Mucus and virus containing tubes were incubated on ice for 2hrs (n=9). Mixture was added to MDCK cells and incubated at 33°C for 2hrs before being removed. The cells were washed and then overlayed like a plaque assay, as described above. Virus growth in the presence of mucus was normalized to a no-mucus control which was used to calculate restoration of infectivity. Statistical significance between groups was compared in GraphPad Prism 10.0.3 via a Mann-Whitney U test. An asterisk indicates statistical significance where *p* ≤ 0.10 (.), *p* ≤ 0.05 (*), *p* ≤ 0.01 (**), *p* ≤ 0.001 (***).

For pseudovirus based assays, mucus was diluted 1:70 in duplicate in PBS in a final volume of 100µL. Mucus was serially diluted twofold in 50µL PBS. 50µL pseudotyped influenza virus was added to each well of a white Nunc™ MicroWell™ 96-well, flat-bottom microplate (pseudovirus was diluted in DMEM to reach a standard titre of 1 × 10^6^ RLU after 48hrs). Mucus and pseudovirus containing plates were incubated on ice for 2hrs to mimic the reverse genetics virus assays. After 2hrs, 50 µL of 1 × 10^5^ HEK 293T cells were added to each well and incubated for a further 48hrs at 37 °C. The cells were then lysed with 50 µL per well of a 1:1 mixture of Bright-Glo^TM^ Luciferase Assay System and 1X PBS. The RLU of the cell lysate was determined using the FLUOstar OMEGA microplate reader. Assays were run in at least three independent biological repeats taken from separate mucus isolations from HNECs.

Microneutralisation assay data was analysed in GraphPad Prism 10.0.3. The reduction of infectivity from the mucus was determined by comparing the RLU in the presence and absence of mucus and expressed as percent inhibition by mucus. Results were normalized to the no-mucus (virus only) condition and to a cell-only control. A neutralisation curve was fit with non-linear regression (inhibitor versus normalised response), which allowed for the calculation of a 50% inhibitory concentration (IC50). The IC50 is the dilution factor of mucus needed to inhibit pseudovirus entry into cells (in RLU) by 50% compared to the control wells. A lower IC50 indicates a higher concentration of mucus is needed to inhibit pseudovirus entry.

## Acknowledgements and Funding

The authors would like to thank Professors Henrik Clausen and Yoshiki Narimatsu of the University of Copenhagen for kindly sharing the HEK293 KO ΔST3GAL1/2/3/4/5/6 ΔST61/2 cells. The authors would also like to thank Professor Nicholas S. Heaton of Duke University for providing the B4GALNT2 plasmid to produce Sda-expressing cells. The authors would like to acknowledge Professor Nigel Temperton of the University of Kent for his contributions to pseudotyped lentivirus development. The authors would like to thank Rebecca Frise of Imperial College London for continued technical support.

Regarding obtaining the viral sequences used in this study, the authors gratefully acknowledge all data contributors, i.e., the Authors and their Originating laboratories responsible for obtaining the specimens, and their Submitting laboratories for generating the genetic sequence and metadata and sharing via the GISAID Initiative, on which this research is based.

HK, KS, JMH, MRO, HH, JY, ADM, YL, JZ, WSB, MI, TPP, SB, J-RS, SJ contribution was supported by BBSRC/DEFRA funded ‘FluTrailMap’ project [BB/Y007298/1], MRC funded FluTrailMap-One Health project [MR/Y03368X/1] and [MR/Y015061/1].

MI, TPP, SB, J-RS, SJ were additionally supported by BBSRC project [APP104179] and Pirbright Institute’s Strategic Programme Grants (ISPGs) [BBS/E/PI/230002A; BBS/E/PI/230001C, BBS/E/PI/230002B] and National Bioscience Research Infrastructure (NBRI) Grants [BBS/E/PI/23NB000, BBS/E/PI/23NB0003].

This work was supported by the Francis Crick Institute (BS & ADS contribution), which receives its core funding from Cancer Research U.K. (CC2127), the U.K. Medical Research Council (CC2127), and Wellcome Trust (CC2127). This work was supported by the Wellcome Trust (218304/Z/19/Z to YL and BS) and UK Research and Innovation (UKRI) under the U.K. government’s Horizon Europe funding guarantee (UKRI3602 to BS).

## Author Contributions

HK, KS, J Yang, ADM, J Yu, and TPP conducted primary virological experiments. MRO, HH, JRS, SB, and SJ prepared cell lines, viruses, and/or proteins in support of these experiments. ADS and BS developed glycan array probes. HK, TPP, YL, and WSB drafted the manuscript. HK, TPP, and WSB designed the study. JH, SMH, MI, YL provided guidance and support to the study design. JM and JZ provided technical support and conducted pilot experiments. All authors reviewed and approved of the manuscript.

## Supplemental Information

**Supplemental Table 1:**
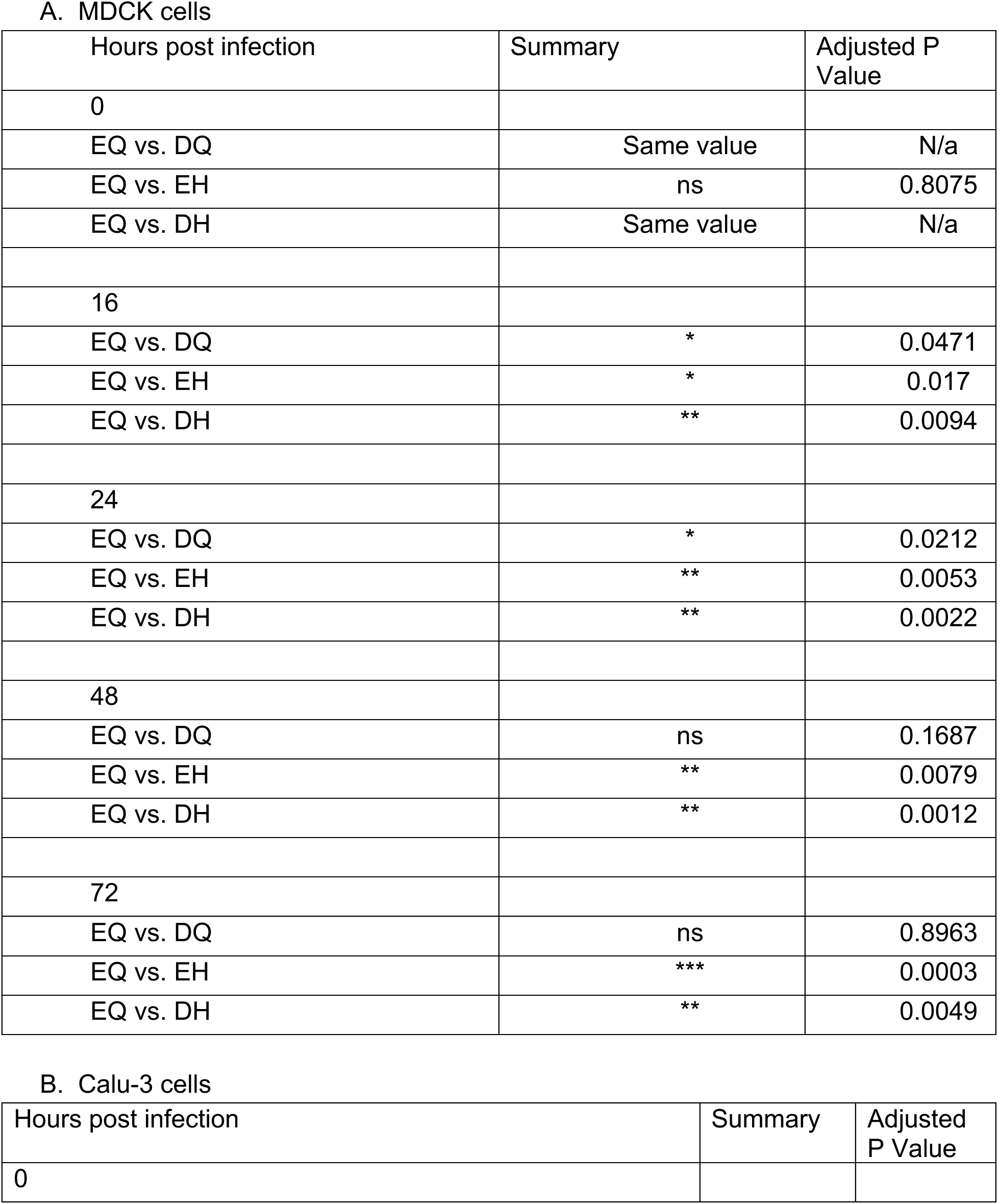
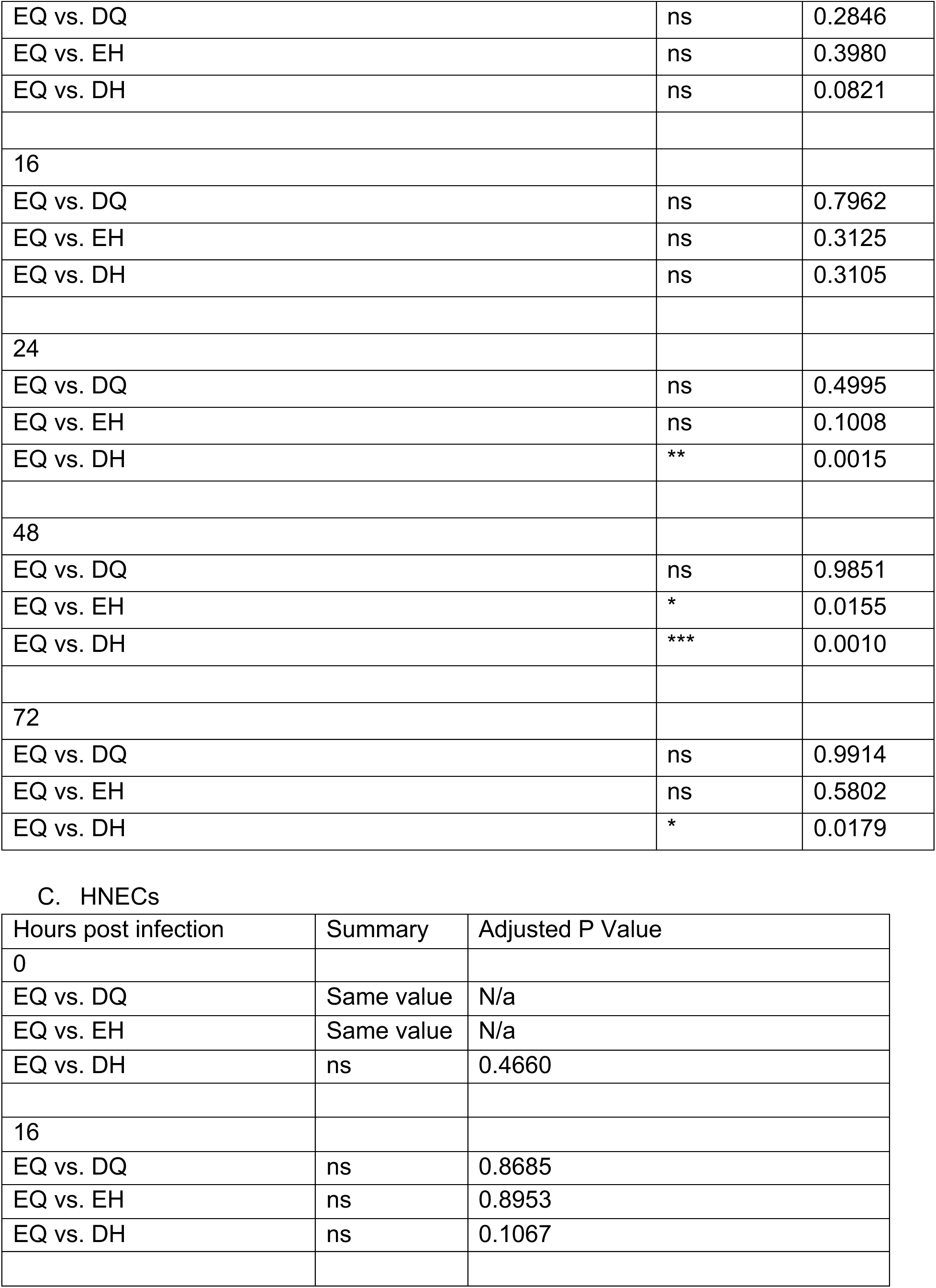
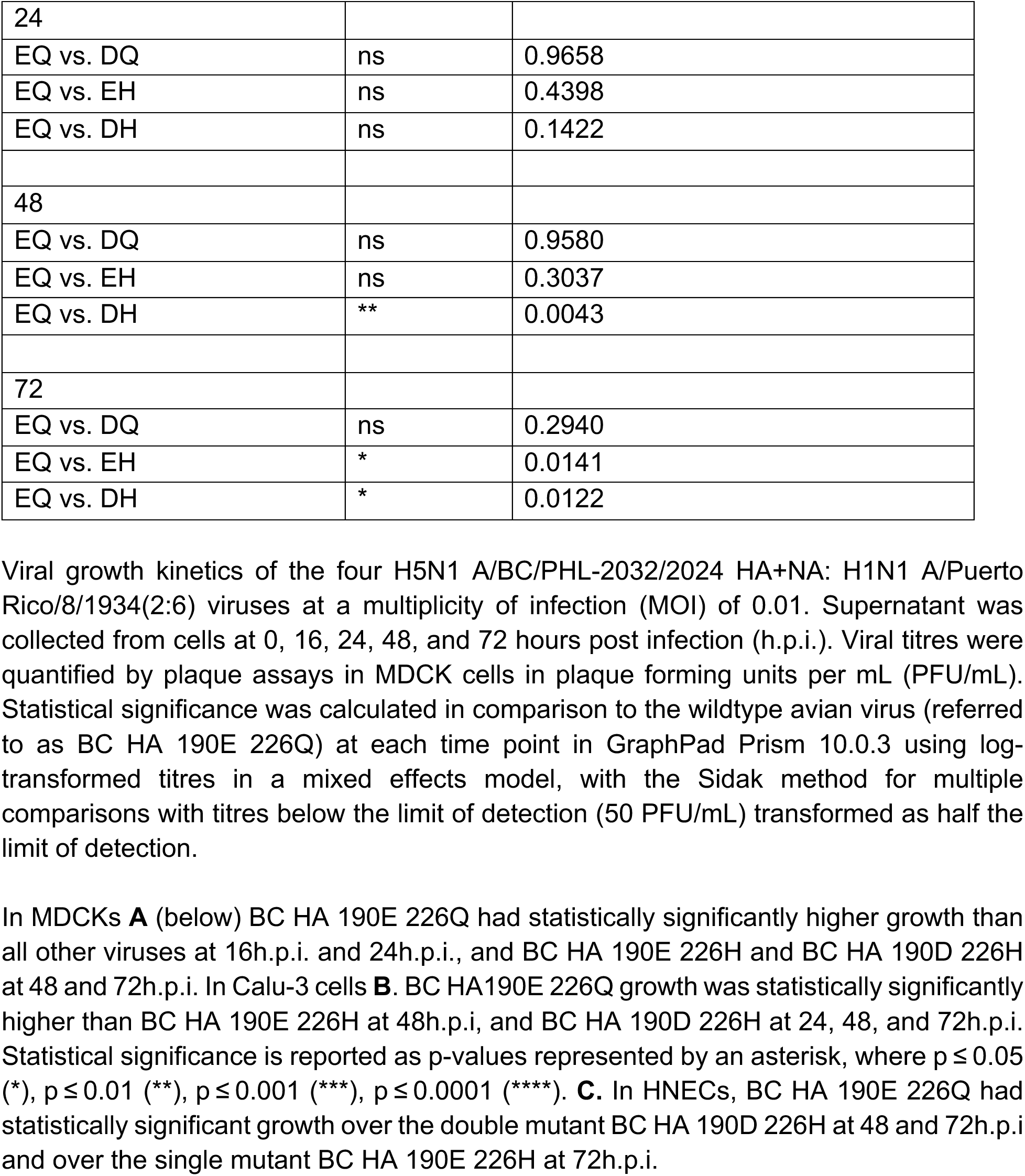
P-values associated with viral replication kinetics curves (Figure 1).

**Supplemental Figure 1:**
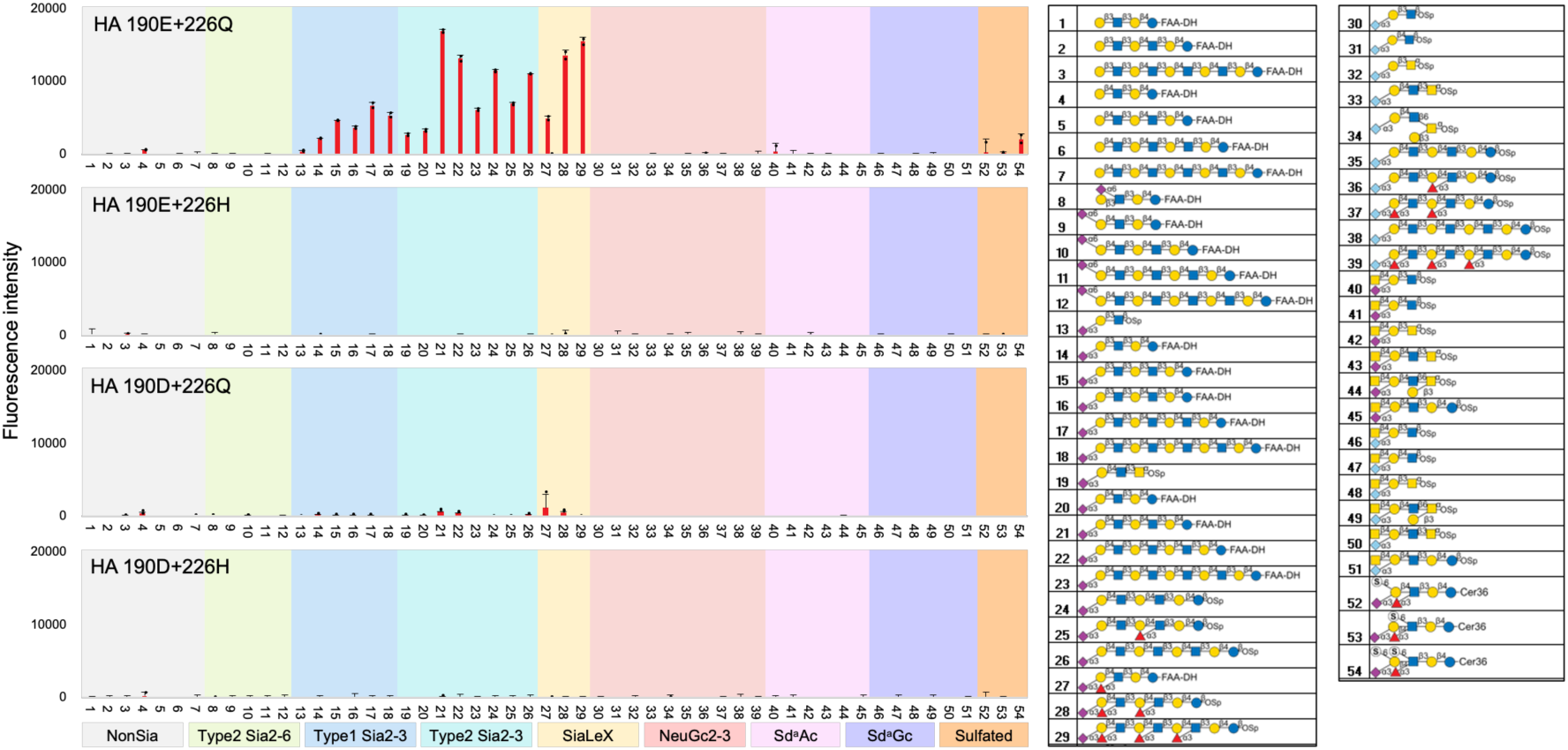
Glycan microarrays with recombinant HA proteins. Glycan microarray binding profiles of the four recombinant BC HA proteins. The 190E 226Q HA showed strong and selective binding to α2-3-linked sialyl glycans, whereas the other three HA proteins showed little or no detectable binding. Data represent the mean fluorescence intensities of duplicate glycan probe spots printed at 5 fmol per spot. Error bars indicate half the difference between the duplicate measurements. The 54 lipid-linked glycan probes are grouped into non-sialylated and sialylated glycans according to their structural features, as indicated by the colour panels. The complete list of glycan probes and binding scores are provided in **Supplementary Data File 1.**

**Supplemental Figure 2:**
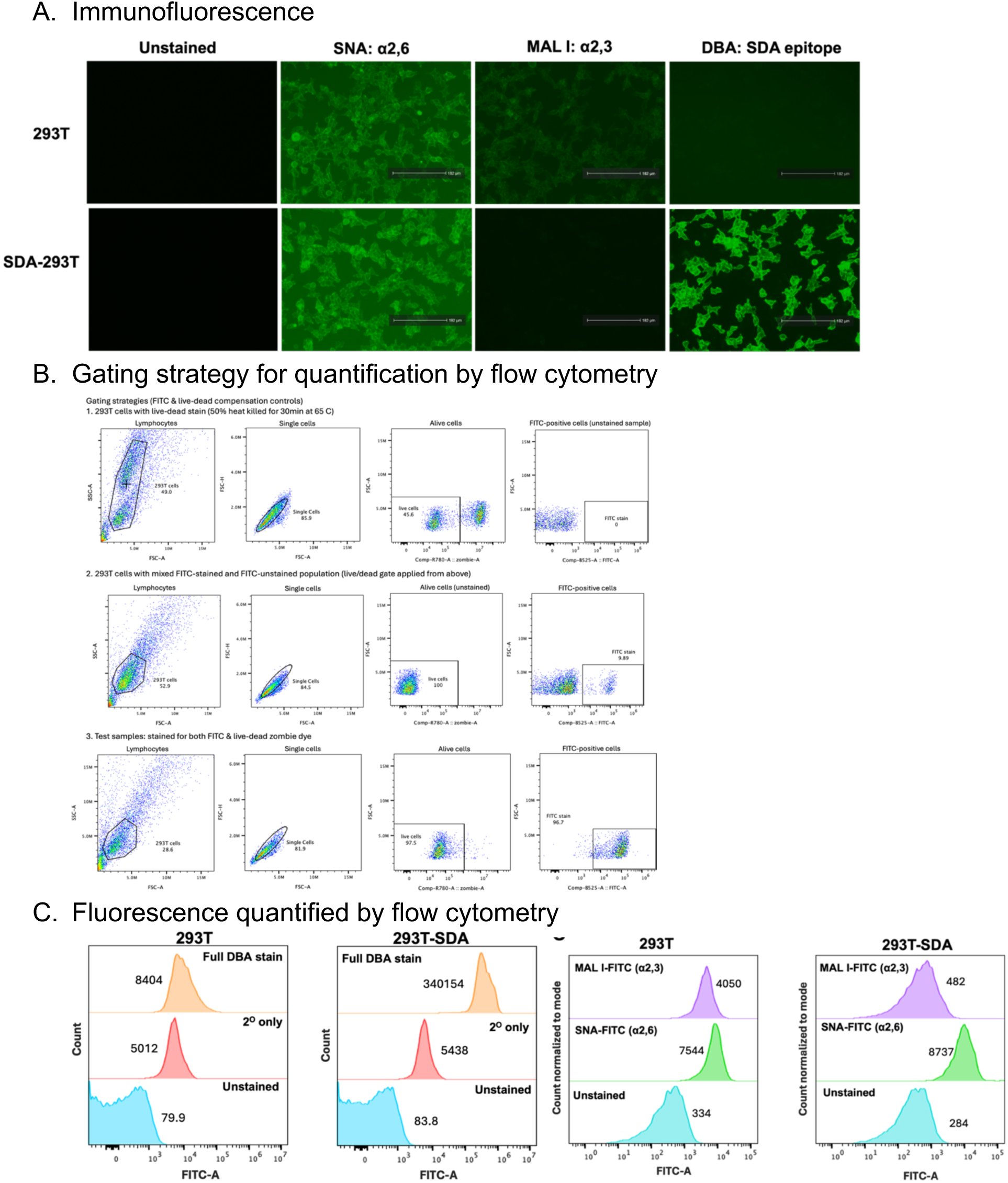
Validation of SDA expressing cell lines. Validation of 293T-Sda cells. **A.** 293T-Sda cells were stained with 20ug/mL lectins conjugated to fluorescein (FITC) in 1X OptiMEM, either MAL I (Vector Laboratories, FL-1311-2), SNA (Vector Laboratories, FL-1301-2), or Dolichos biflorus lectin (DBA; binds the Sda epitope, Sigma-Aldrich). Cells were incubated with the lectins for 1hr at 4°C, then washed once with 200μl OptiMEM. DBA lectin was conjugated to biotin, so after 1hr these wells were washed and stained with streptavidin-conjugated Alexa Fluor™ 488nm (Invitrogen, S11223) for 1hr at 4°C. Cells were stored in PBS and visualised with an EVOS M3000 Imaging Auto fluorescent microscope. The 293T-Sda cells showed reduced MAL-I staining compared to the wildtype cells, suggesting that Sda was successfully blocking a2,3-linked sialic acids. **B.** Cell surface expression for 293T-Sda cells was also quantified by flow cytometry. Cells were stained with Zombie-NIR fixable viability dye (BioLegend, 423105) to determine live vs dead populations. Cells were then fixed as in the immunofluorescence studies above. Cells were stained with lectins as in **A**. Events were then acquired on the NovoCyte Penteon flow cytometer (Aligent Technologies). A minimum of 10,000 events was collected per condition, and results were gated to exclude doublets and dead cells. **C**. The Sda cells had reduced MAL-I staining with unaffected SNA staining, suggesting that α2,3-linked sialic acids were successfully modified by Sda.

**Supplemental Table 2:**
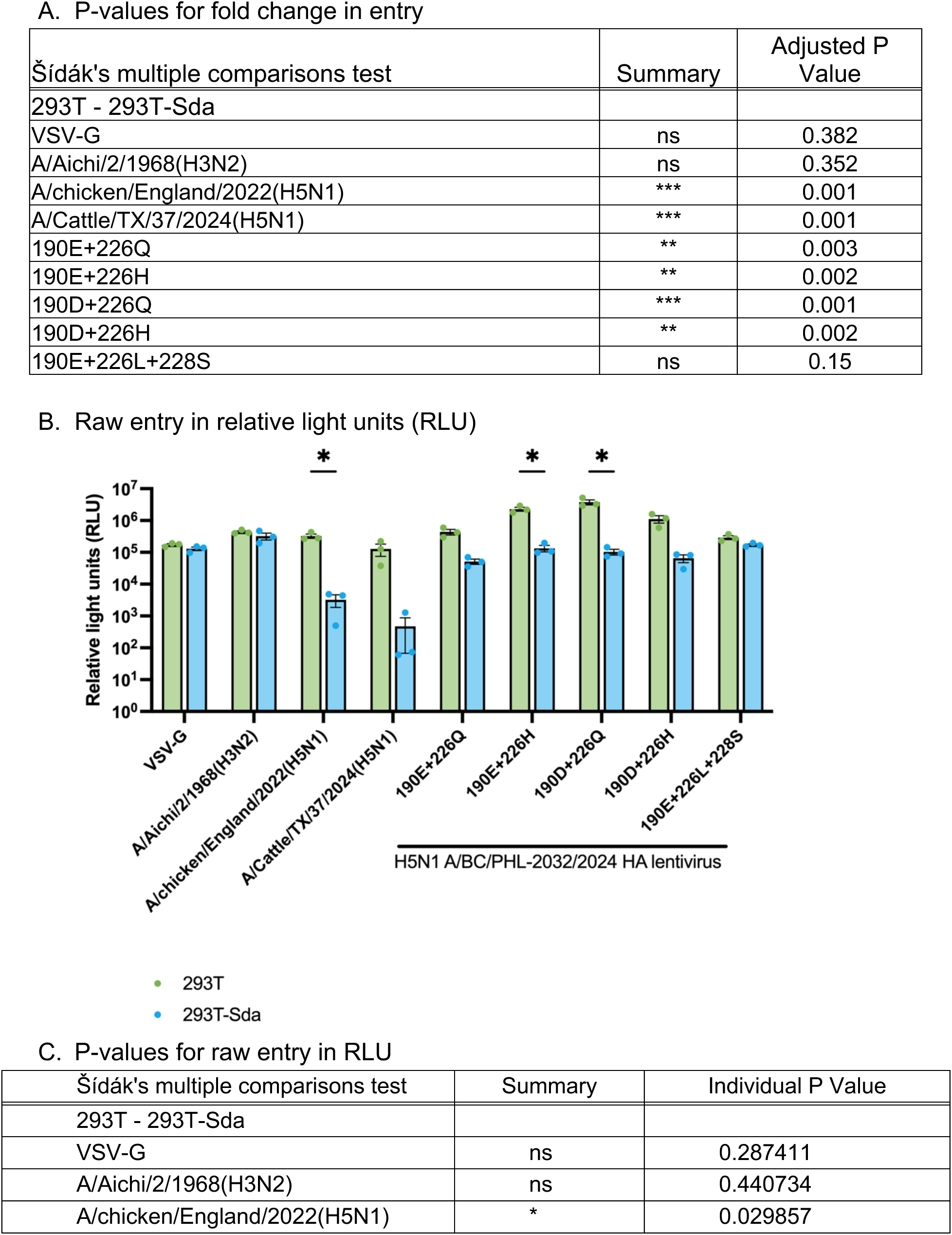
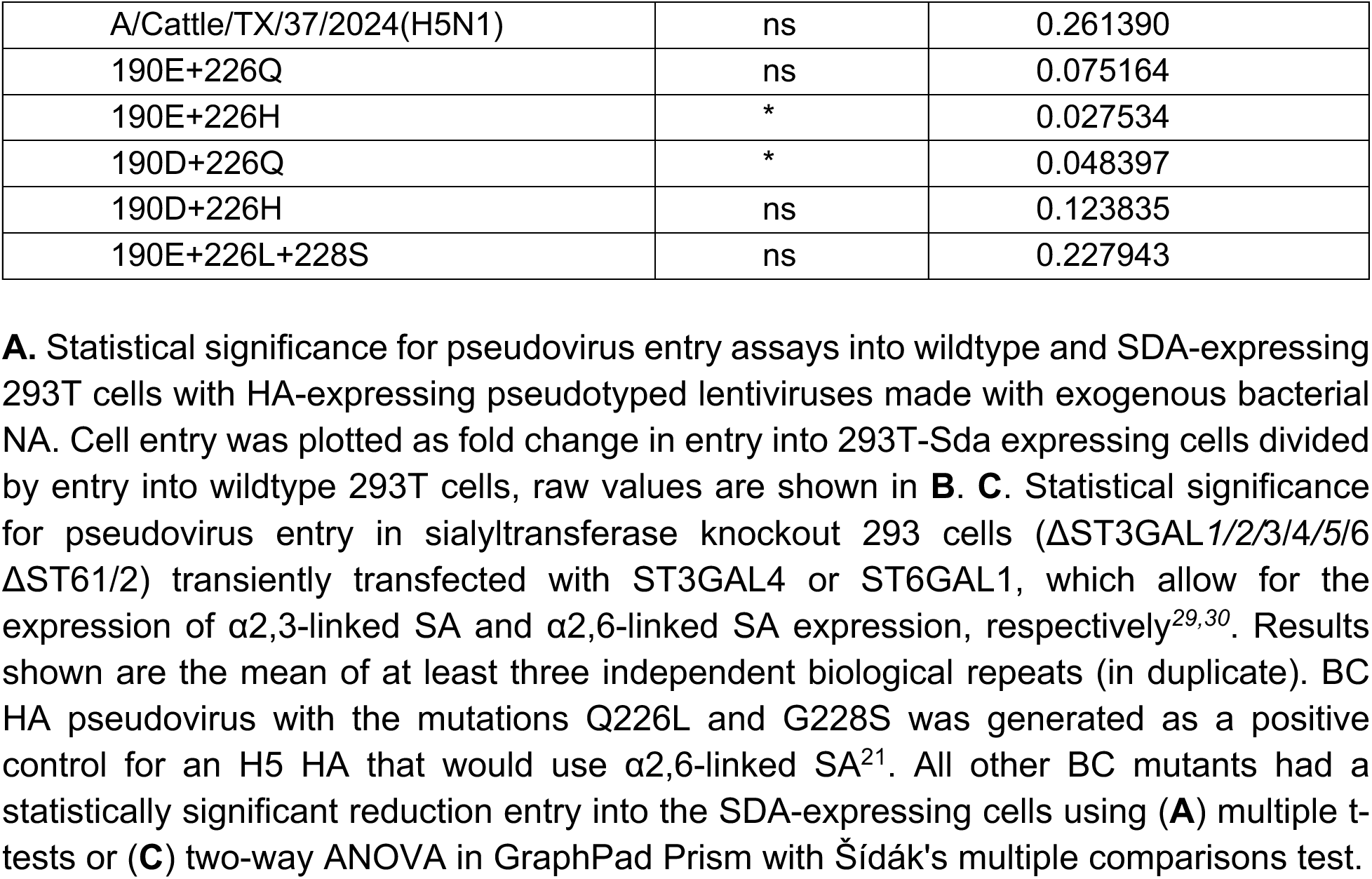
Pseudovirus entry into wildtype vs SDA-expressing 293T cells (Figure 2)

**Supplemental Table 3:**
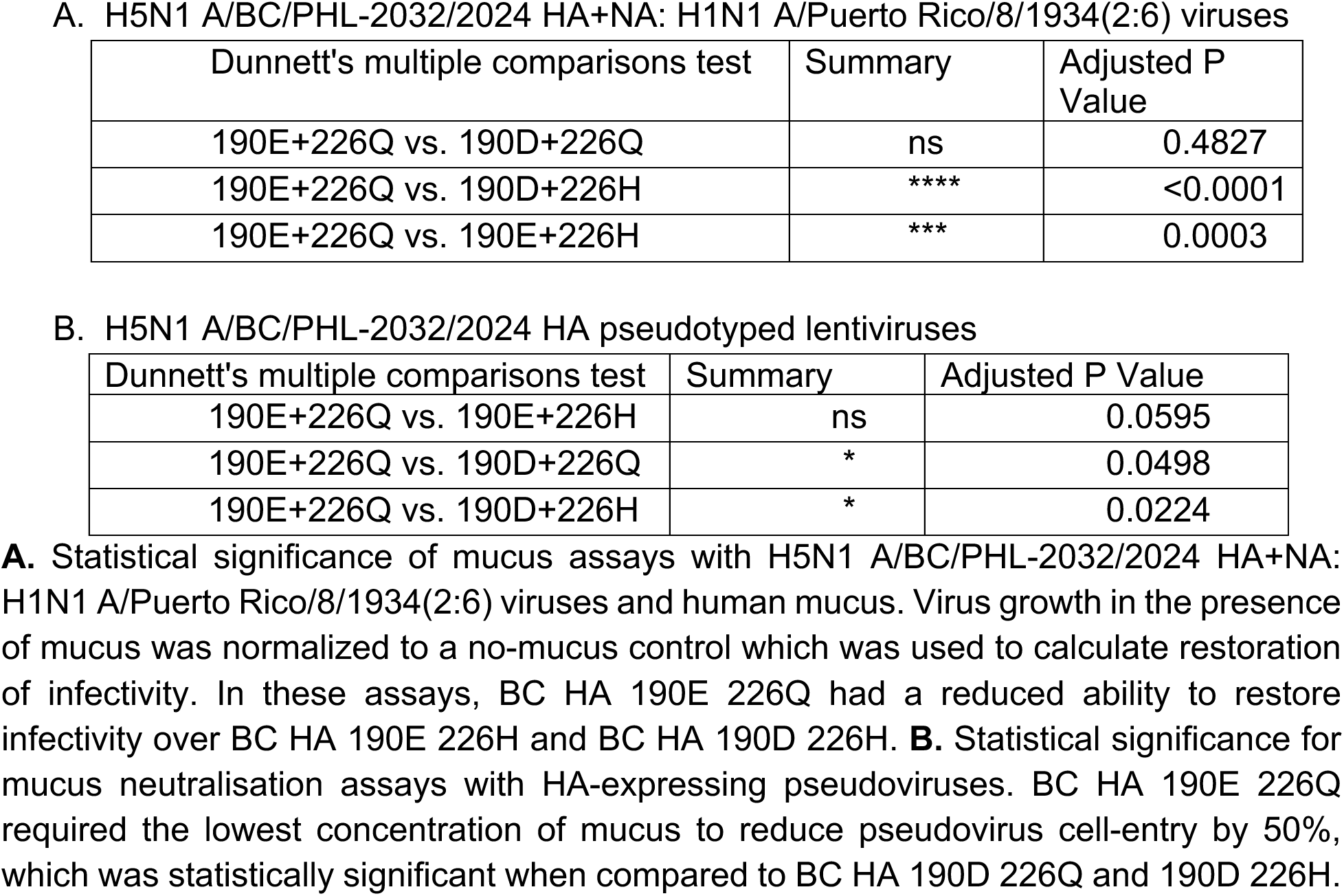
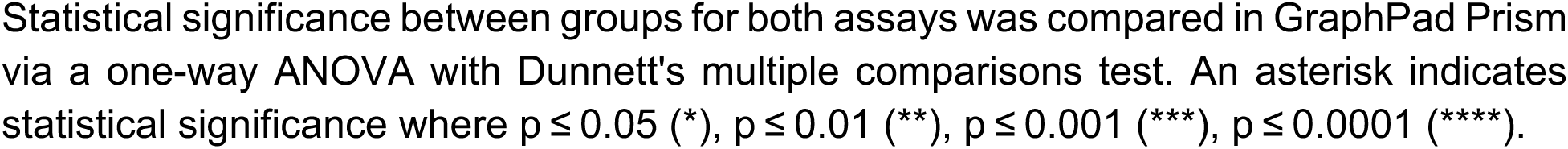
Group comparison of mucus assays (Figure 3)

**Supplemental Table 4:**
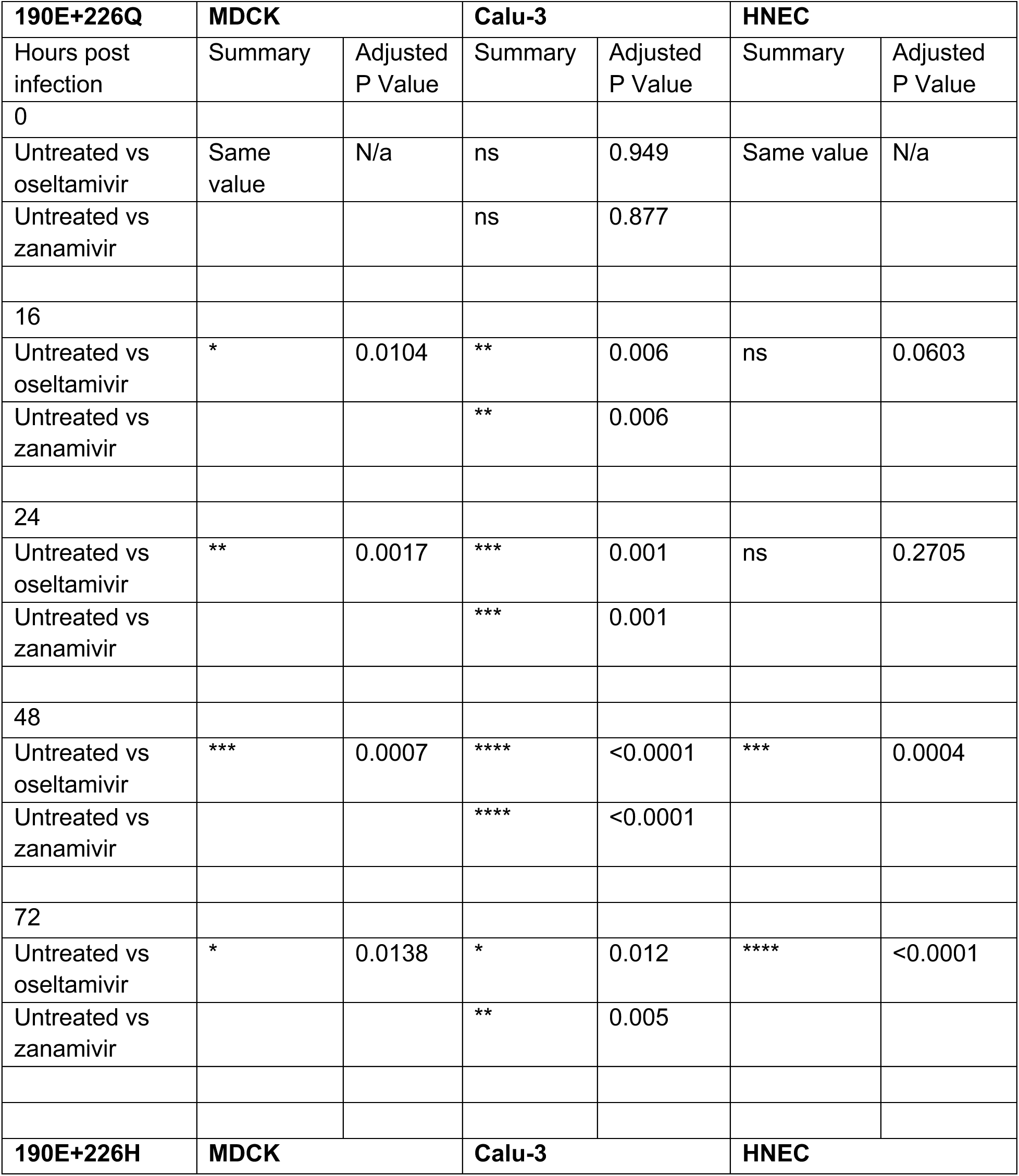
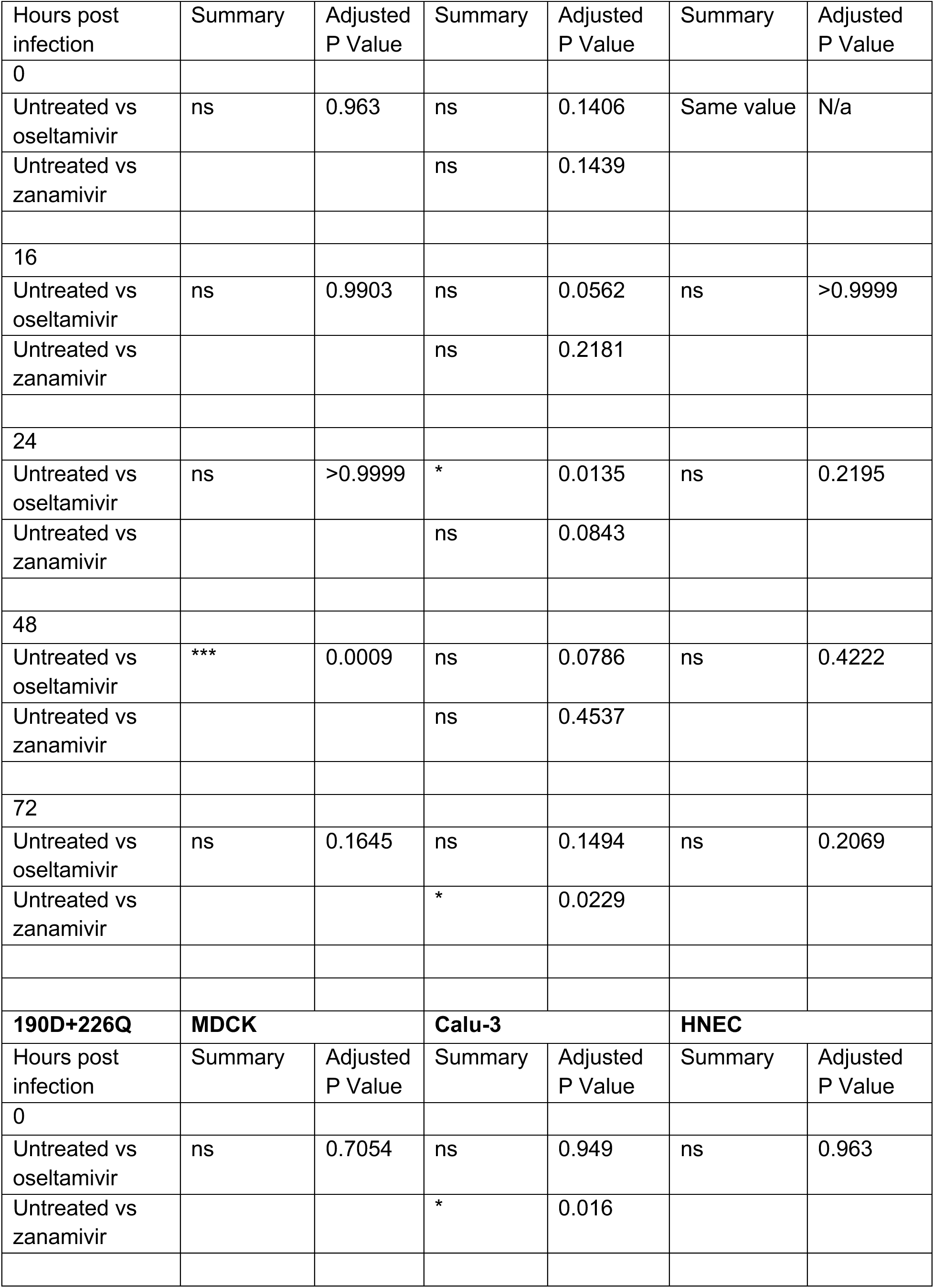
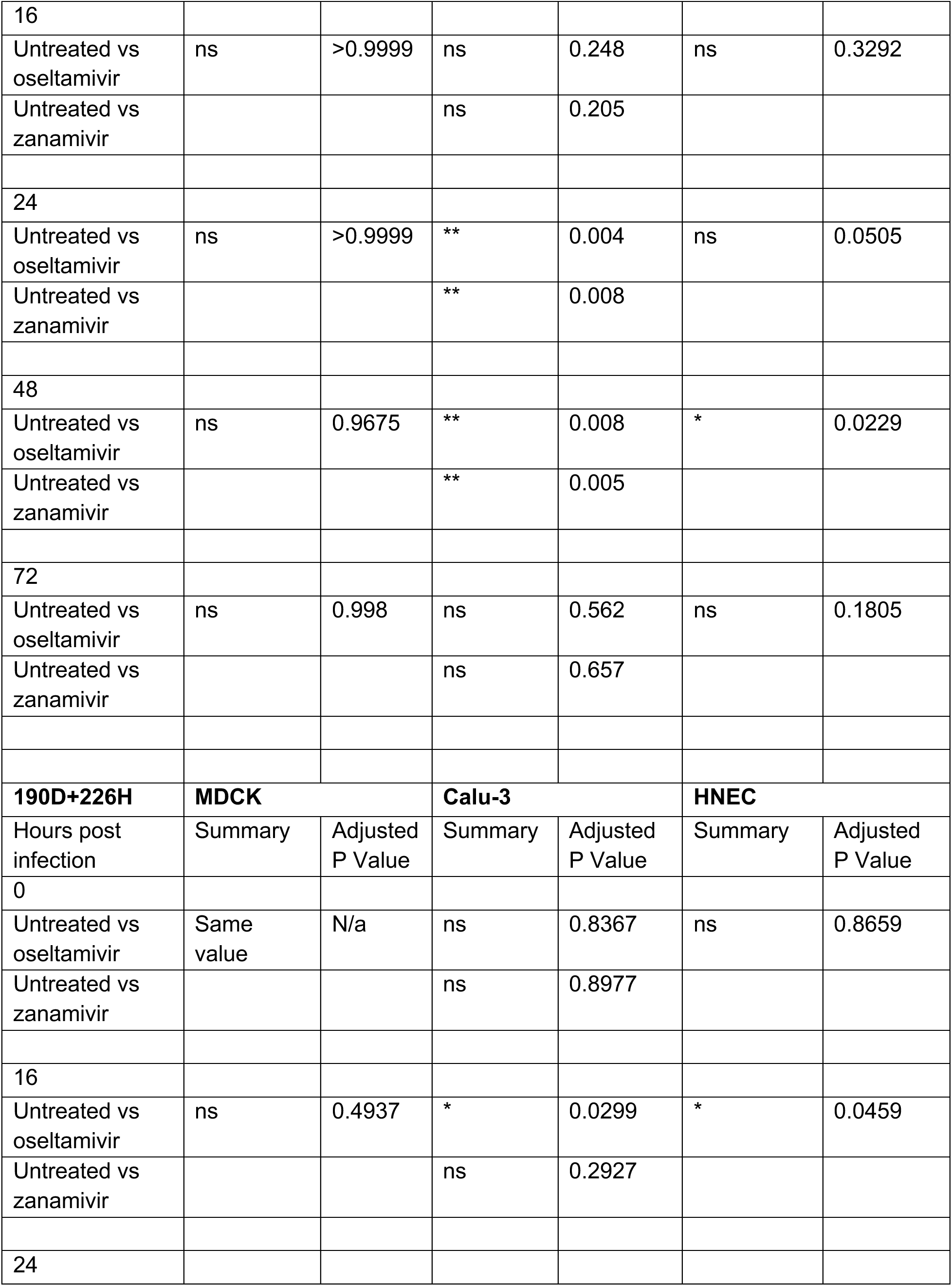
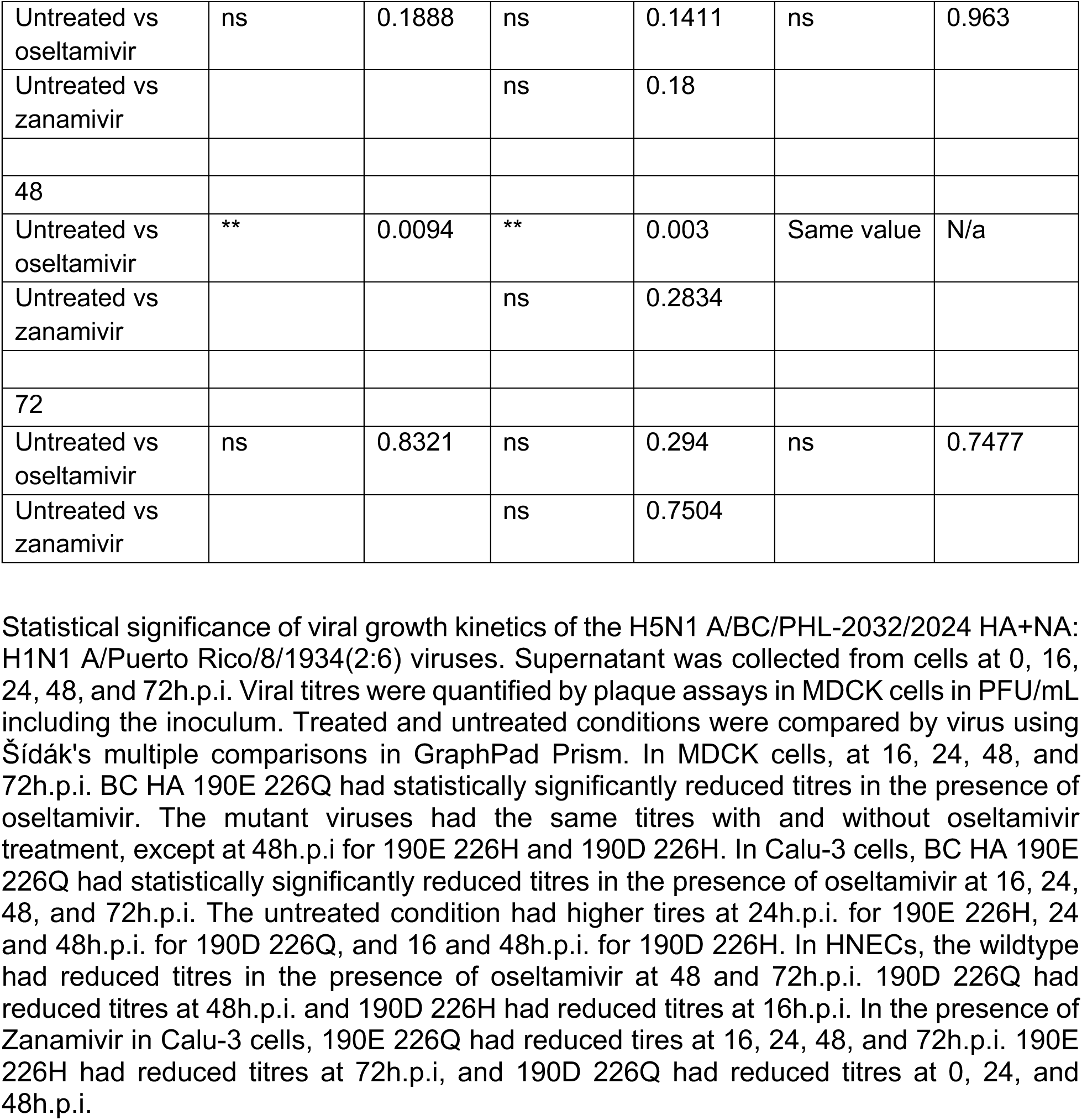
Šídák’s multiple comparisons test associated with viral replication kinetics curves in the presence of oseltamivir or zanamivir (Figure 4).

**Supplemental Figure 3:**
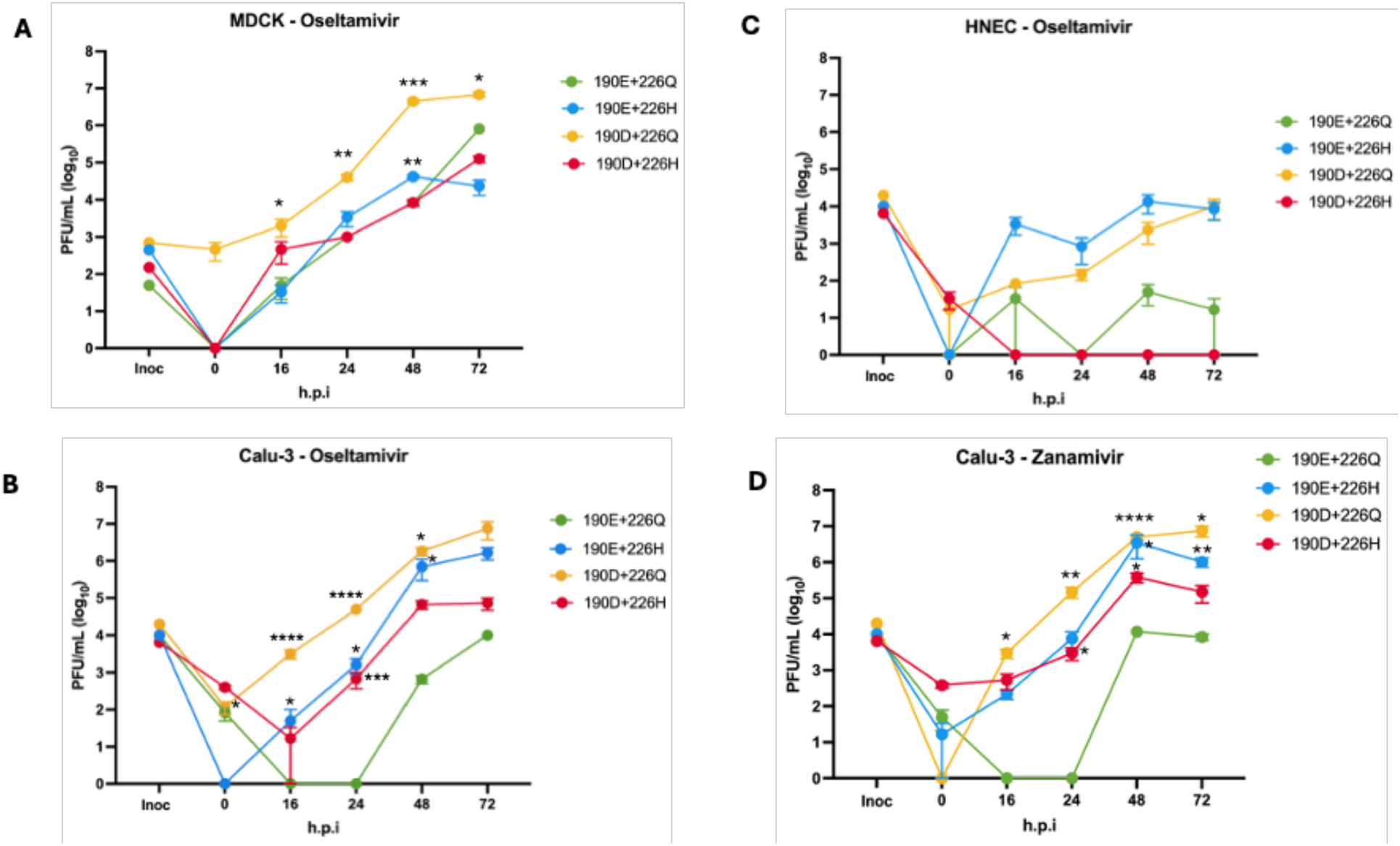
Viral replication kinetics curves in the presence of oseltamivir or zanamivir. Data from Figure 4 was then compared relative to wildtype virus (this data is replotted from Figure 4). Viral growth kinetics of the H5N1 A/BC/PHL-2032/2024 HA+NA: H1N1 A/Puerto Rico/8/1934(2:6) viruses in the presence of oseltamivir. Supernatant was collected from cells at 0, 16, 24, 48, and 72h.p.i. Viral titres were quantified by plaque assays in MDCK cells in PFU/mL including the inoculum, labelled Inoc using viruses at MOI 0.01. Each point represents the mean with of three biological replicates (n=3) with the standard deviation. **A**. In MDCK cells, at 0, 16, 24, 48, and 72h.p.i. BC HA 190D 226Q had statistically significantly higher titres than the wildtype. At 48h.p.i, BC HA 190E 226H had statistically significantly higher titres than the wildtype. The titres of the wildtype and double mutant were not statistically different at any time point. **B**. In Calu-3 cells, BC HA 190D 226Q had statistically significantly higher titres at 16, 24 and 48h.p.i. BC HA 190E 226H had higher viral titres at 16, 24, and 48h.p.i. than the wildtype. BC HA 190D 226H had statistically significantly higher titres than the wildtype at 24h.p.i. **C**. In HNECs, the wildtype and mutant titres were not statistically different at any point. **D**. In the presence of Zanamivir in Calu-3 cells, BC HA 190D 226Q had higher titres than the wildtype at 16, 24, 48, and 72h.p.i. BC HA 190E 226H had higher titres than the wildtype virus at 48 and 72h.p.i. BC HA 190D 226H had higher titres than the wildtype at 24 and 48h.p.i. Statistical significance for viral replicative fitness was calculated as previously. BC HA 190E 226Q and BC HA 190D 226Q expressing pseudoviruses had statistically significant decreases in entry without oseltamivir. Statistical significance is reported as p-values (**Supplemental Table 5**) represented by an asterisk, where p ≤ 0.05 (*), p ≤ 0.01 (**), p ≤ 0.001 (***).

**Supplemental Table 5:**
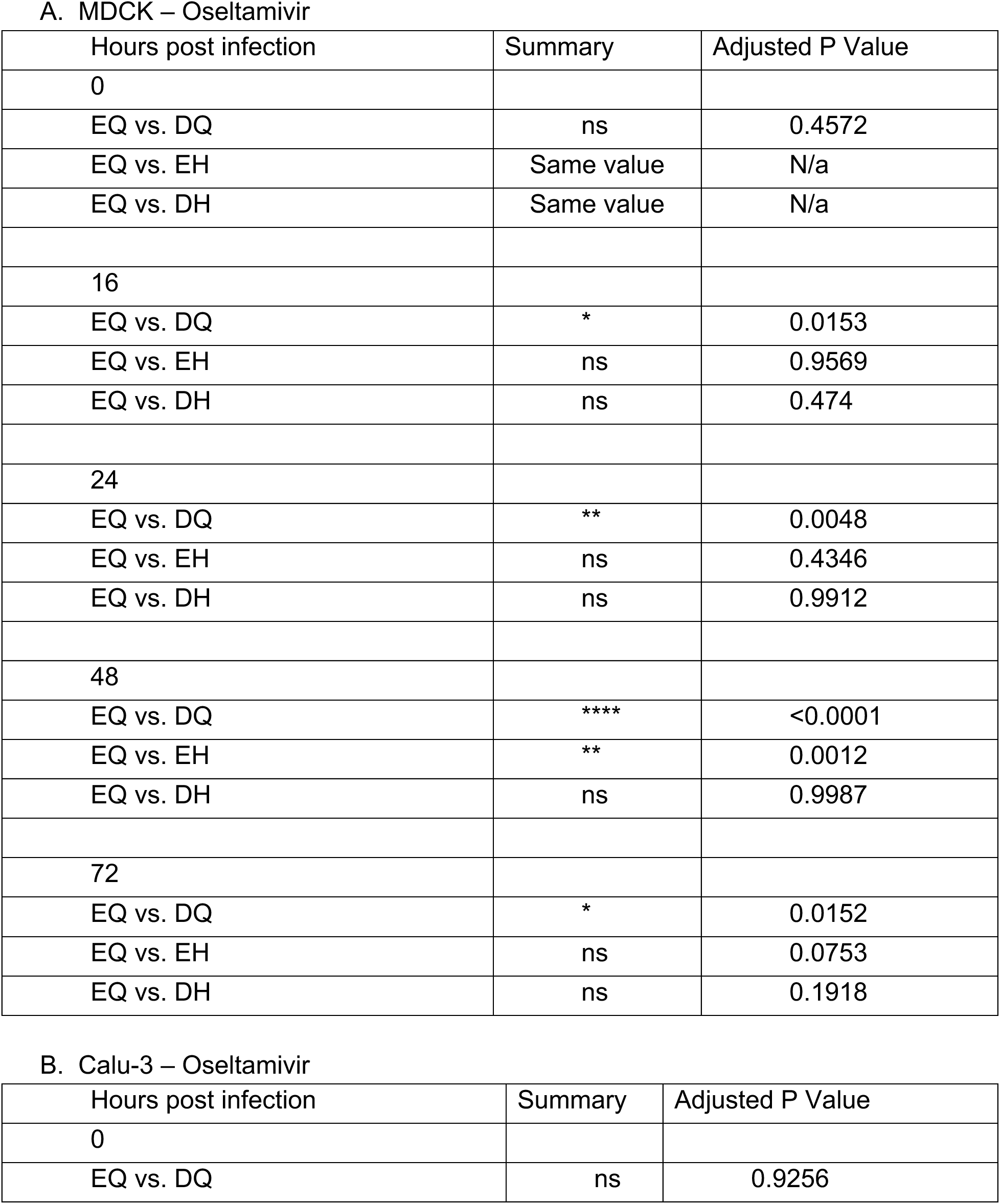
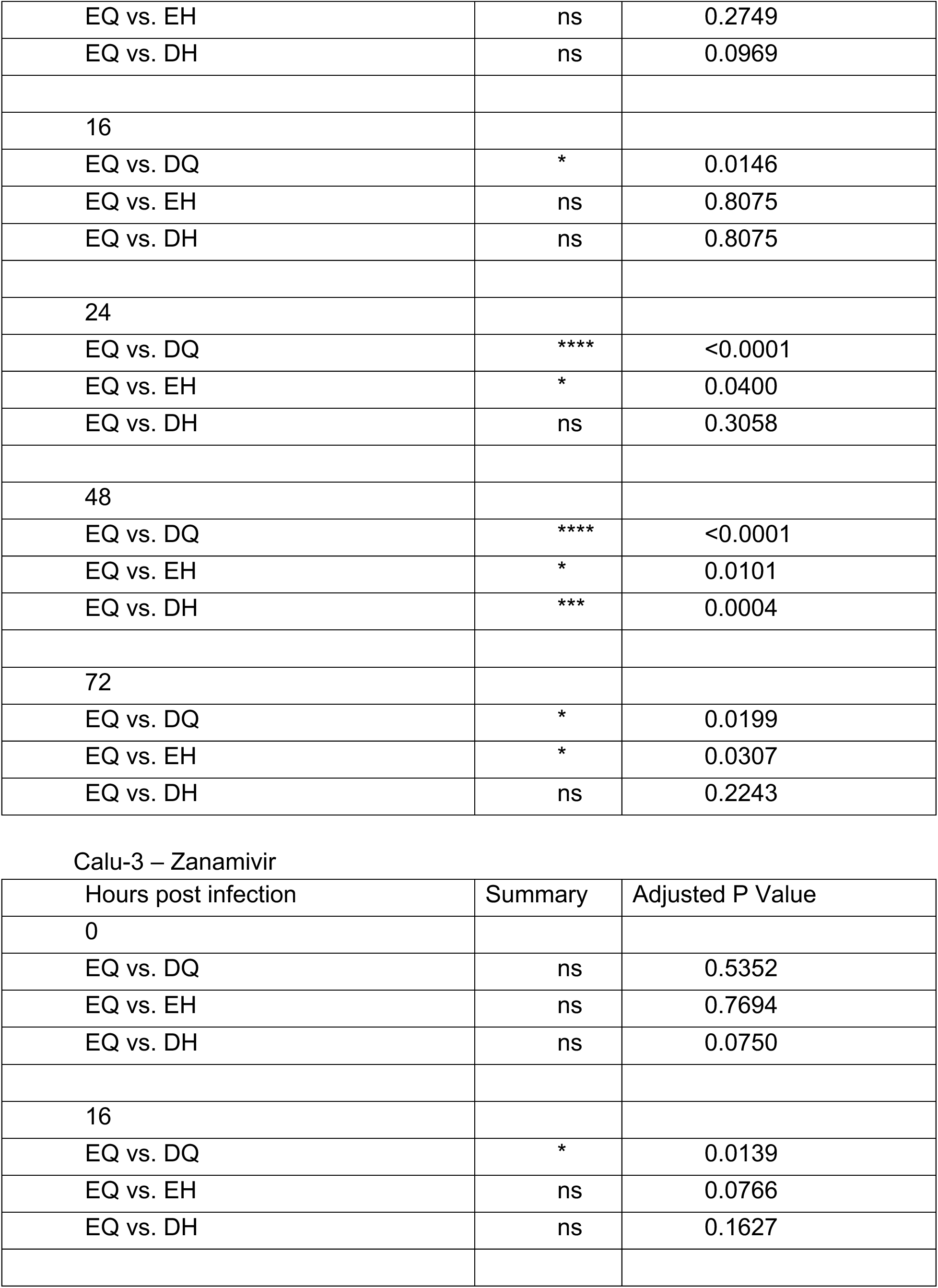
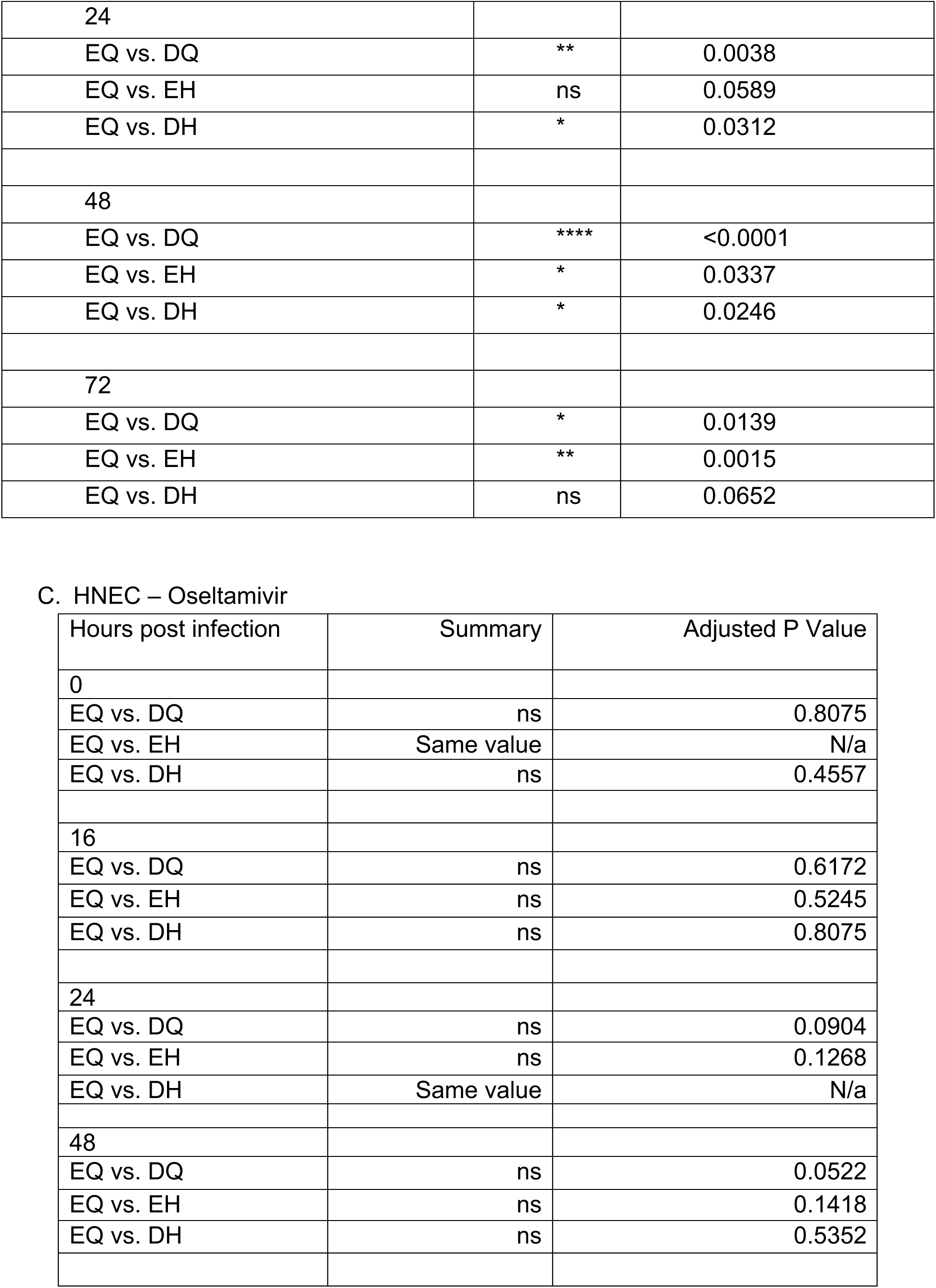
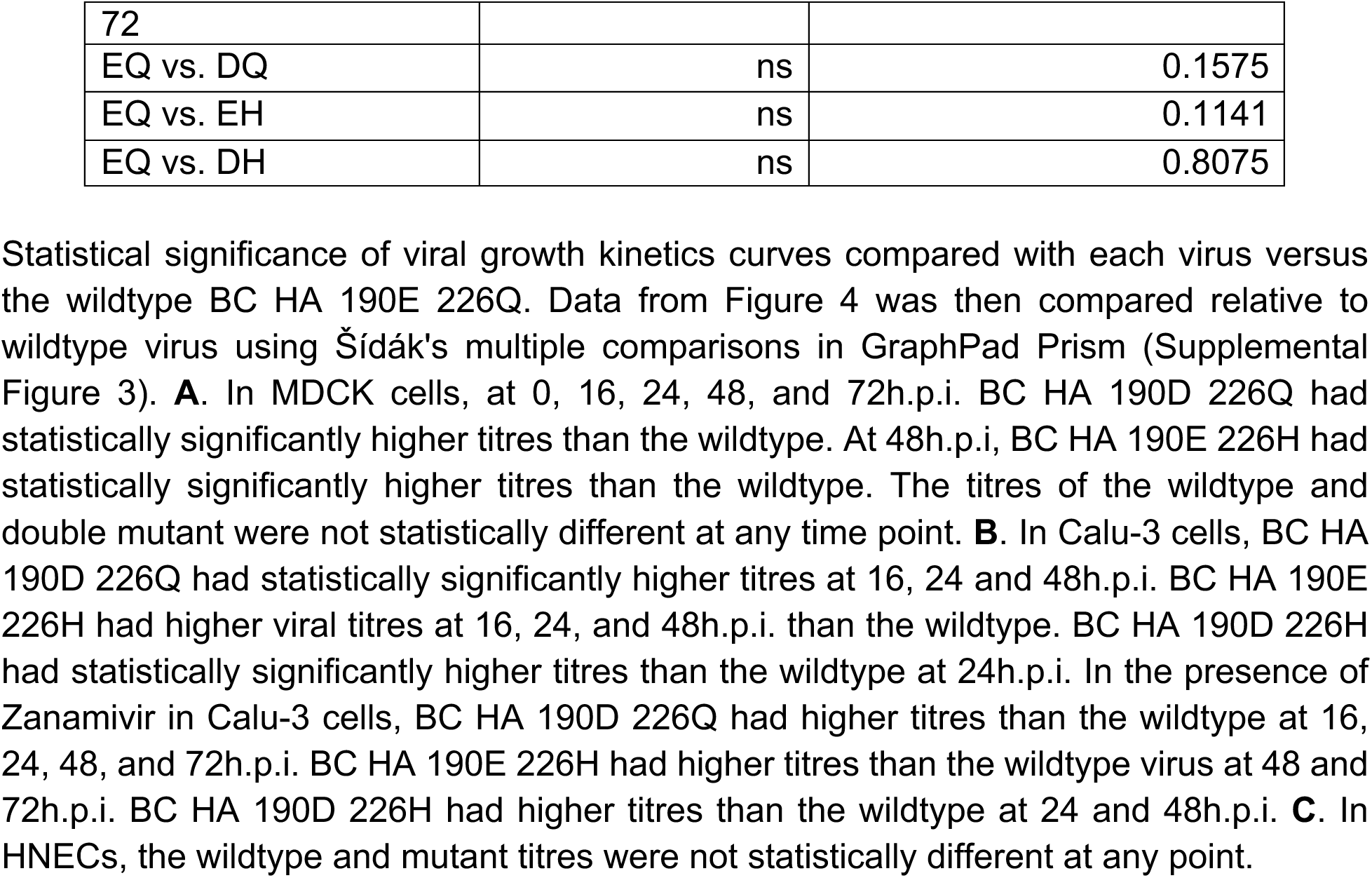
Šídák’s multiple comparisons test associated with viral replication kinetics curves in the presence of oseltamivir or zanamivir (Supplemental Figure 3).

